# Non-Stationary Dynamic Mode Decomposition

**DOI:** 10.1101/2023.08.08.552333

**Authors:** John Ferré, Ariel Rokem, Elizabeth A. Buffalo, J. Nathan Kutz, Adrienne Fairhall

## Abstract

Many physical processes display complex high-dimensional time-varying behavior, from global weather patterns to brain activity. An outstanding challenge is to express high dimensional data in terms of a dynamical model that reveals their spatiotemporal structure. Dynamic Mode Decomposition is a means to achieve this goal, allowing the identification of key spatiotemporal modes through the diagonalization of a finite dimensional approximation of the Koopman operator. However, DMD methods apply best to time-translationally invariant or stationary data, while in many typical cases, dynamics vary across time and conditions. To capture this temporal evolution, we developed a method, Non-Stationary Dynamic Mode Decomposition (NS-DMD), that generalizes DMD by fitting global modulations of drifting spatiotemporal modes. This method accurately predicts the temporal evolution of modes in simulations and recovers previously known results from simpler methods. To demonstrate its properties, the method is applied to multi-channel recordings from an awake behaving non-human primate performing a cognitive task.

## I. INTRODUCTION

Data-driven models of spatio-temporal systems are critical to understanding the evolution dynamics of natural systems and have become especially valuable given the increasing prevalence of large-scale measurements across all scientific disciplines. Many methods have been introduced to derive approximate dynamical models from data in domains such as fluid flows [39], climate systems [45, 1] and brain activity [53, 65, 25]. However, in many data-driven approaches and algorithms, the data are assumed to be stationary when fitting the data. The stationarity assumption is violated in many datasets of interest, thus limiting potential model accuracy and forecasting capabilities. Deriving non-stationary generalizations of data-driven modeling is an area of active interest (e.g. [22, 32, 35]). We add to this effort by proposing a new method, Non-Stationary Dynamic Mode Decomposition (NS-DMD), which explicitly approximates the non-stationarity of the data while simultaneously constructing a low dimensional linear DMD approximation of multi-variate time-series.

NS-DMD builds on Dynamic Mode Decomposition (DMD) [51, 31], a systematic and unbiased method to reduce high-dimensional time-series data to a set of spatiotemporal modes. DMD approximates the Koopman operator [49], a linear infinite dimensional operator whose eigendecomposition models the observables that describe a finite dimensional, potentially non-linear, dynamical system [30, 5, 56]. In short, DMD approximates the data **x**(t) as

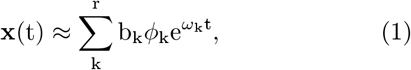

where the *ϕ*_k_ are the DMD modes, *ω* are the DMD eigenvalues and b_k_ determines the weight of each mode. The limitation of such an approach is the assumption of stationarity of the data. Simulated datasets of particular interest (Fig. 1(A)) are poorly reconstructed with stationary approaches (Fig. 1(B)) due to the spatial mixture of time-varying modes with time-varying amplitudes in the data. NS-DMD improves upon DMD by including time-dependence of the modes:

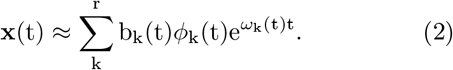

The additional time dependence in NS-DMD allows for accurate reconstruction of the underlying data (Fig. 1(C)). Further details on the NS-DMD method are found in Sec. II-B.

**FIGURE 1:**
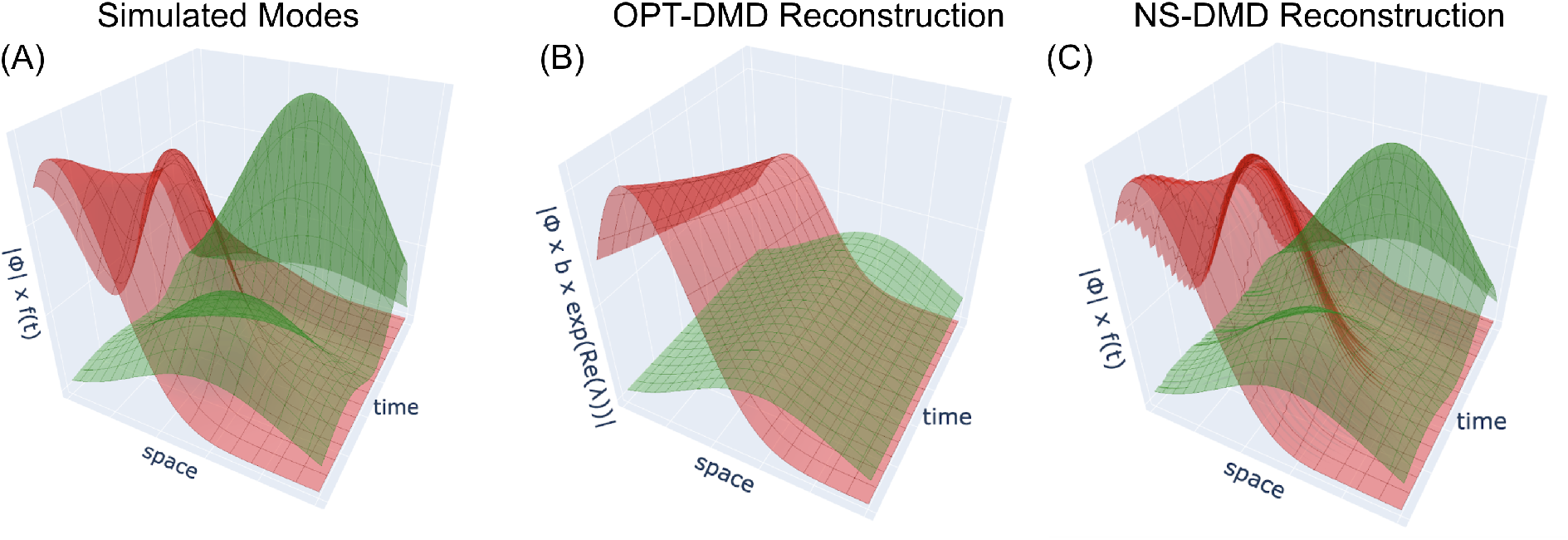
(A) The spatial distribution |*ϕ*_k_| and global amplitude f_k_(t) of a simulated dataset, originating from the form 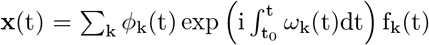. The frequencies *ω*(t) are 20 Hz (red mode) and 30 Hz (green mode). The spatial modes mix together at the same time with varying amplitudes. The red modes spatial distribution fluctuates while the green mode turns on and off with irregular amplitudes. OPT-DMD fails to recover the original modes (B). NS-DMD samples the data with OPT-DMD during different time intervals, where dynamics are combined to find a subset of modes with modulating amplitude to reconstruct the data. NS-DMD recovers the original modes (C). Further exploration of NS-DMD and OPT-DMD on non-stationary datasets is shown in Fig. 3.

Many approaches exist to fit non-stationary systems [15, 6, 59, 16, 12, 17, 8, 23, 22, 57, 28, 35, 32, 9, 41], but the approaches do not find representations in the form of Eq. 2. Related work is discussed in Sec. IV-A.

As with many other methods, NS-DMD assumes that the data are stationary in small time windows. We further assume that the data contain a low dimensional set of spectral components which may vary slowly with respect to their frequency. However, in contrast to previous methods, NS-DMD subsequently takes advantage of machine learning methods to associate modes across time windows, while systematically eliminating overfit and redundant modes. This allows us to detect global modulations of spatiotemporal modes that gradually drift across time.

We validate NS-DMD on synthetic data from several non-stationary systems. We then demonstrate its practical utility by analyzing multi-channel neuroscience time-series data. NS-DMD is able to recapitulate results found using other more traditional time-series analysis methods, but also identifies non-stationary modes in these time-series data. We further demonstrate the utility by applying NS-DMD to sea surface temperature data, where NS-DMD recovers seasonal effects along with modes specific to the El Niño phenomena. Taken together, the novel findings and the connection to previous methods demonstrate the promise of NS-DMD for the analysis of non-stationary data.

## II. METHODS

### NOTATION

We follow the notation in [29]. Scalars are denoted by lowercase letters (s), vectors by bold lowercase letters (**v**), matrices by bold capital letters (**M**), and tensors of third order by calligraphic bold letters (*𝒯*). In summary:

- v_i_ denotes the ith entry of **v**;
- m_ij_ denotes element (i, j) in **M**;
- t_ijk_ denotes element (i, j, k) in *𝒯* ;
- **m**_i:_ and **m**_:j_ denote the ith row and jth column of **M**;
- More compactly, **m**_j_ ≡ **m**_:j_, denotes the jth column of **M**;
- **t**_ij:_, **t**_i:k_, and **t**_:jk_ denote the vectors given by the corresponding free dimension of *𝒯* ;
- **T**_i::_, **T**_:j:_, and **T**_::k_ denote the matrices given by the corresponding free dimensions of *𝒯* ;
- More compactly, **T**_k_ ≡**T**_::k_ denotes the kth frontal slice;
- The nth element in a sequence is denoted by a superscript in parenthesis; e.g. **M**^(n)^ is the nth matrix **M**.

#### A. DYNAMIC MODE DECOMPOSITION

Dynamic Mode Decomposition (DMD) [31] forms the backbone of NS-DMD. DMD approximates a lowdimensional representation of the data **X** in terms of linearly (exponential) evolving spatial modes (Eq. 1). That is, at fixed frequencies given by Im(*ω*_k_), there are spatial modes *ϕ*_k_ with loadings b_k_ that exponentially grow or decay. DMD is thought to combine the strengths of singular value decomposition across space with Fourier transforms across time.

There have been many improvements made to DMD since the original algorithm was introduced in 2008 [51], including a number of regression techniques for estimating the best fit linear dynamics [61, 2, 50, 63, 32, 44]. We build NS-DMD upon Optimized DMD (OPT-DMD) [2], which estimates the DMD modes and eigenvalues by using a variable projection optimization scheme

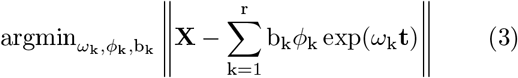

where a rank r approximation is estimated. Optimized DMD iterates to a solution of this non-convex problem by using variable projection [18]. To improve convergence capabilities, often the exact DMD algorithm can be used as a seed for the initialization of the DMD algorithm. The OPT-DMD framework has been found to be the most robust algorithm to noise [2], providing an unbiased estimate of the DMD modes and eigenvalues for real data.

#### B. NON-STATIONARY DYNAMIC MODE DECOMPOSITION

For a data matrix **X** ∈ ℝ ^*N* × *M*^, DMD aims to accurately represent the data with a low dimensional set of K spatiotemporal modes 𝒮 ∈ ℝ^*N* × *K* × *M*^ as given by Eq. (3). When the governing processes vary in time, the set of spatiotemporal modes that best describe the data at one point in time may not describe the data well at another point in time. Furthermore, nonlinear dynamical systems may be better described locally by different linear approximations in different regions of phase space. We postulate that there are common dynamical modes that recur throughout an extended dataset. The goal of NS-DMD is to find these recurring modes and weight their amplitude with time-dependent functions **F** ∈ ℝ^*K* × *M*^ that characterize the time-varying contribution of each mode to a reconstruction of the data. Thus, NS-DMD seeks to approximate the data at any snapshot t_j_ with

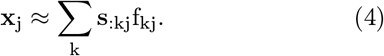

To approximate 𝒮, a common approach [22, 32, 41, 4] is to split the data **X** of length M into W short overlapping windows of length P < M, 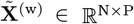, defined by the corresponding set of time points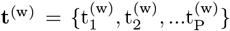. The processes governing the dynamical systems are assumed to be approximately stationary in each window, an assumption that is valid depending on the size of the window. If the size is too large, then the stationarity assumption is likely false. If the size is too small, there is a lack of statistical sampling to find reasonable solutions that may not generalize well across time. For example, in the limit of two snapshots, it is unlikely for modes to generalize to any another snapshots.

The data of each window are extracted with OPTDMD [2], an iterative algorithm that finds r modes per window, where r is a chosen hyperparameter. For the first sampled window 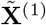, OPT-DMD is executed without initial conditions or with educated guesses; OPT-DMD can automatically determine an initial guess if needed [2, 31]. For sampling windows w > 1, OPT-DMD is initialized with the normalized eigenvalues **Λ**^(w−1)^*/* |**Λ**^(w−1)^|from the previous window, a process that favors spatiotemporal smoothness.

Having determined modes **Φ**^(w)^ ∈ ℂ^N×r^, **Λ**^(w)^ ∈ ℂ^r×r^, and **b**^(w)^ ∈ ℝ^r^, local to every window w, the goal is to identify a set of modes that apply across the full M length dataset. A visual description of this process is shown in Fig. 2 (a). Formally, similar spatiotemporal modes are grouped into K groups across consecutive windows 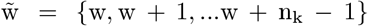, a process defined and explained in Sec. II-B2. We write this as 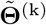, where each 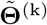 contains a variable number n_k_ of modes 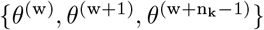, and **Θ** is a placeholder for **Φ, Λ, b**, and **t**. To find time dependent quantities 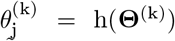 at time t_j_, the function h averages 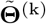 during overlapping windows and extrapolates 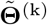 outside the range of **t**^(k)^:

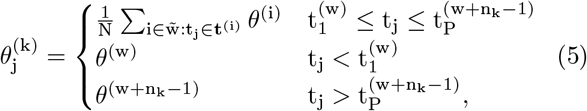

where N refers to the number of summed terms. We use a notation where 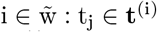 indicates all elements 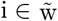 such that t_j_ ∈ **t**^(i)^. A visualization of the function h is shown in Fig. 2 (b). The application of h leads to 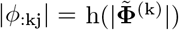 and 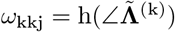. The angular part of **Φ**^(w)^ defines the phase of every channel at the start of the respective window. To find ∠*ϕ*_:kj_, the phases are aligned to the start of the full dataset. For a mode k and window w_i_, the phase is 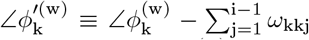, and the phase at any time is 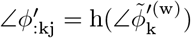. To gain greater temporal smoothness, one can apply a moving average to the time dependent modes. The k time dependent spatiotemporal modes form the matrices **Φ**_j_ ∈ ℂ^*N* ×*K*^ and ∠**Λ**_j_ ∈ ℂ^*K* ×*K*^ for every time point t_j_. With the time dependent modes, the spatiotemporal modes 𝒮 ∈ ℝ^*N* ×*K* ×*M*^ are:

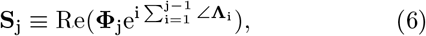

where the real part of each mode recovers the complex conjugate pairs in the definition of **S**_j_.

**FIGURE 2:**
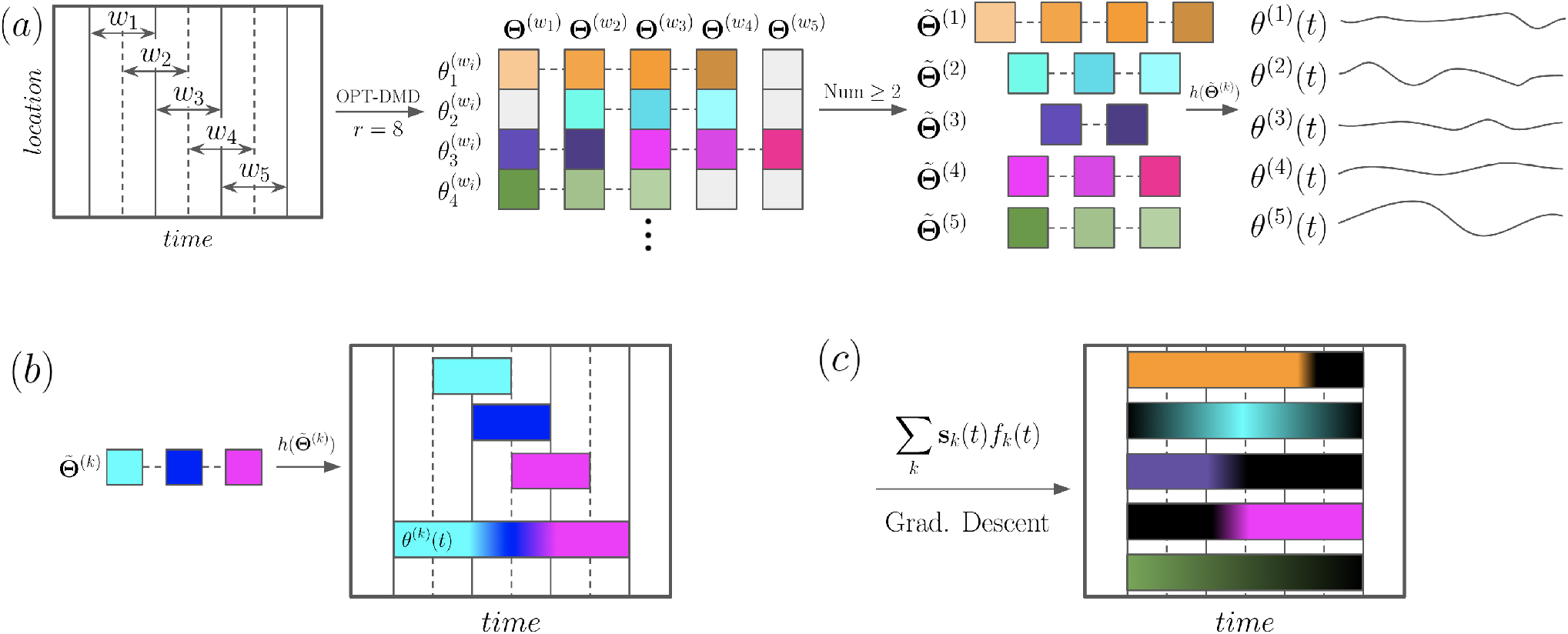
Flowchart of the NS-DMD algorithm. In (a), a data matrix (first panel) is subdivided into windows. The number, size, and stride of the windows are hyperparameters. For each window, OPT-DMD is computed with r modes. The modes are visualized as squares in the second panel, where 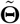 is a placeholder for the OPT-DMD modes: **Φ** and **Λ**. Modes are deemed similar to each other based on the procedure in Sec. II-B2; similar modes are connected by dashed lines and have similar colors. Different shades indicate that the modes may not be exactly the same. Groups of minimum 2 (another hyperparameter) similar modes (third panel) are transformed into continuous modes across time in the fourth panel via the function h 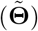. The function h 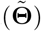 is visualized in (b). Similar, but slightly different modes, are indicated by slightly different colors. For example, the frequency could be 10Hz in window 2, 10.2Hz in window 3, and 10.4Hz in window 4. For overlapping windows, modes are averaged and extended outside the range of the windows. Following the example, the frequency would be 10.2Hz before window 3 begins, 10.1Hz during the overlap of windows 2 and 3, 10.3Hz during the overlap of windows 3 and 4, and 10.4Hz for the remainder. The modes *θ*^(k)^(t) comprise the spatiotemporal modes 𝒮 (see text). Lastly, temporal modulations f_k_(t) of each mode **S**_k_ are found with gradient descent. f_k_(t) is visualized in (c), where the colored bars indicate when each mode well describes the data. f_k_(t) is flexible and can find gradual changes to the modes (e.g. the green and light blue mode). Other times, the modes turn on/off rapidly (e.g. the orange, purple, and pink modes). Lastly, the timing is flexible, indicated by the modes not necessarily turning on/off exactly at the dashed lines.

𝒮 is weighted with an unknown temporal modulation **F** ∈ℝ^*K* ×*M*^. A visualization of **F** is shown in Fig. 2(C). The time-series is reconstructed with

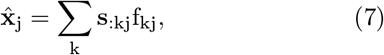

where the vectors 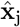 form the estimated data matrix 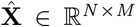. To solve for **F**, we use gradient descent (Sec. II-B1). An alternate, “exact” method is proposed in the supplementary text (Sec. S1-A), which aims to directly solve Eq. 7. However, this method tends to be noisier than gradient descent.

##### 1) Fitting Time-Varying Modes with Gradient Descent

Constraints are needed before applying gradient descent to find **F**. First, **F** is non-negative; assemblies are either “on” with some variable amplitude, or they are “off.” Second, a sparsity constraint is added since 𝒮 may contain redundant modes. Lastly, **F** is continuous since we assume that modes turn on or off on the timescale defined by the sampling rate 1*/*sr = Δt.

The following loss function satisfies these constraints on **F**:

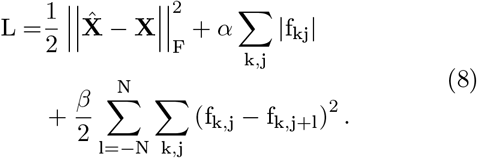

The first term is a standard least-squares loss term on the reconstruction of the data given **F**. The *α* term enforces sparsity in the solutions of **F**. The absolute value will be dropped since **F** is forced to be non-negative after every iteration of gradient descent. Lastly, the *β* term enforces continuity across time; *β* controls the degree of smoothing, and the smoothing timescale is controlled by N. For simplicity, *β* is fixed for each l, although in principle *β* can fluctuate.

The gradient of **F** for each mode k and snapshot t_j_ is

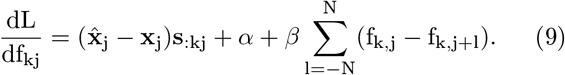

Finally, this allows us to compute the gradient descent for an iteration i > 1:

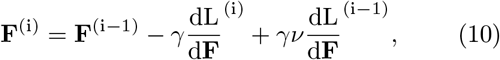

where *γ* is the learning rate and *ν* is the momentum [47].

The initial guess of f_kj_ is found by setting 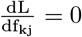 and solving for *α* = 0 and **f**_k′≠k, l ≠0_ = 0:

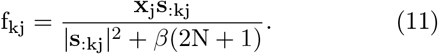

To remove noise, the initial guess is lowpass filtered.

After each iteration, all negative values of **F** are set to 0, and 2N + 1 consecutive values are averaged to further control the smoothness of **F. F** is reflected at the boundaries. It is possible for edge artifacts to occur when **F** > 0, so a minimum of N values should be trimmed at the boundaries.

After running gradient descent, the average amplitude of each mode is typically smaller than the true value due to *α, β*, and the averaging step. The amplitude, defined as **a** ∈ ℝ^*K*^, of each mode is adjusted with the least squares algorithm: **x**_j_ = Σ_k_ **s**_:kj_a_k_f_kj_. The amplitude **a** is absorbed into **F**.

##### 2) Feature Selection

Typically, while estimating **S**, one finds a large number of redundant modes, even with the sparsity constraint in gradient descent. We turn to feature selection to subselect modes.

To combine redundant spatiotemporal modes, the similarity of pairs **S**_i_ and **S**_j≠i_ is determined. The frequencies, spatial amplitudes, and spatial phases are all needed to be similar for two modes to be defined as similar. The difference in frequencies f = ∠λ/(2π) must be within a desired threshold: |f_i_ − f_*j*≠*i*_| < thresh. The cosine similarity 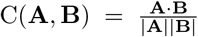 between spatial amplitudes |*ϕ*| must be above a desired threshold: C(|*ϕ*_i_|, |*ϕ*_j≠ i_|) > thresh. The spatial phases ∠*ϕ* need to first be aligned since they are referenced to their window’s initial t_0_. To align, we define Δ∠*ϕ* ≡ (∠*ϕ*_i_ + 2*π*f_i_Δt*/*2) − (*ϕ*_j≠i_ − 2*π*f_j≠i_Δt*/*2), where Δt = t_0,j≠i_ − t_0,i_. Due to periodicity, all Δ∠*ϕ* are shifted to within −*π* to *π*. The spatial phase is similar if 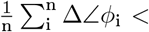 thresh. The solutions **S**_i_ and **S**_j≠i_ are considered similar (i.e., redundant) if all three threshold checks are valid.

The first redundant set of modes are found from the parity of spatiotemporal modes. If the data is sufficiently approximated by sines and cosines, then OPT-DMD returns pairs of solutions **S** with opposite signs: 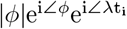 and 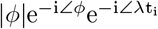. The real part of **S** is compared between pairs, where one mode is removed per pair.

Next, the modes from consecutive windows are compared. Since OPT-DMD is computed on each window using the eigenvalues of the previous window as an initial guess, the modes typically remain similar across time unless the assembly drastically changes. In some cases, the frequency or spatial distribution of modes may fluctuate over time. A method that is very sensitive to such changes would generate additional new modes. Our modeling goal is to have a single mode describe the fluctuation, so the method needs to be flexible when the frequency or spatial distribution drifts in time. This is later tested with great success in Sec. III-A2. If modes are similar across time, we stitch them together according to Eq. 6, as described in Sec. II-B.

Next, if desired, one can retain only groups of modes that have more than a user-defined minimum number of consecutive similar windows. This enforces that 𝒮 is somewhat consistent across time. Then, the reconstruction error is calculated for each mode independently, defined as the cosine distance between **X** and the reconstruction 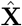. The cosine distance is chosen since it does not depend on the amplitude of each mode. A user-defined number of modes or any modes with a reconstruction error above a user-defined threshold are retained. This drastically reduces the number of modes to ones that generally reconstruct the data well.

After initially reducing the number of modes, standard feature selection methods [3] are used to find the subset of the remaining modes that best reconstruct the data. The basic feature selection algorithms of [46] have been implemented. These methods start with either none or the entire set of modes and add/subtract one mode at a time while checking the reconstruction error. We first run gradient descent (Sec. II-B1) while adding/removing each mode independently. Then, the mode that decreased the cosine distance the least is removed. The process is repeated until a final, user selected number of modes have been added/removed. The best sub-selection of modes can be chosen from an “elbow” curve of overall cosine distance as a function of number of modes.

##### 3) Sampling from a Broad Frequency Range

A dataset may contain a large number of modes at many different frequencies. Running OPT-DMD with many modes can be slow and inaccurate if some frequency bands have smaller amplitudes. To compensate, an additional step to NS-DMD is to bandpass over many different frequency ranges. Initial guesses of frequencies should lie within the bandpass ranges, and OPT-DMD is computed for each window in each band. Only solutions where the frequency is within the bandpassed range are included. All modes from all bandpassed ranges are combined in the feature selection step. Gradient descent is ran on the original, non-bandpassed data.

Either a type I Chebyshev filter of order 5 and a pass-band gain of 1 or a Butterworth filter of order 5 is used to bandpass filter the data. Due to edge artifacts at the temporal extremes of the data, it is recommended to bandpass for a longer time range than of interest and trim the excess timepoints.

To determine the accuracy of the reconstruction during sequential feature selection methods, it’s best to calculate the reconstruction similarity for each frequency band:

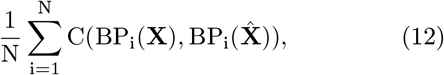

where BP_i_(**A**) is a function that bandpasses the data **A** to the ith frequency range. This method of calculating the reconstruction similarity does not preferentially bias toward modes that occur with a comparatively large amplitude.

##### 4) Non-Stationary Dynamic Mode Decomposition Algorithm

###### Algorithm 1 NS-DMD

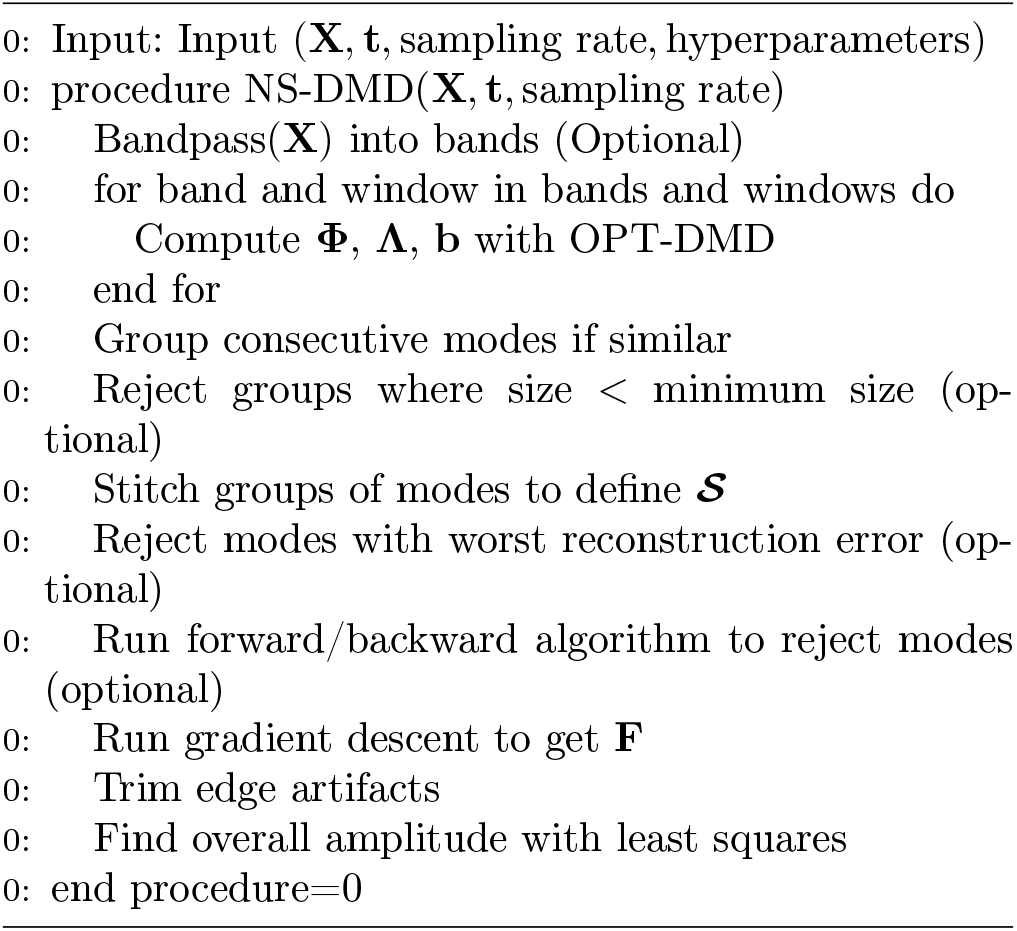

We conclude this section by summarizing the full method. An algorithmic version is given in 1.

- Step 1 (optional): If the data has a large number of modes or includes modes with a much smaller amplitude, we bandpass the data into many different small bands. To evaluate the necessity, we suggest analyzing the power spectral density.
- Step 2: Run OPT-DMD for every window of interest and for every frequency band if applicable.
- Step 3: Find the similarity of consecutive modes.
- Step 4 (optional): Keep only solutions that are similar for a user defined number of windows.
- Step 5: define **𝒮** based on the similarity of consecutive modes. Optionally include temporal lags for each recording location.
- Step 6 (optional): Initially reduce the number of modes by finding the reconstruction error of each mode independently.
- Step 7 (optional): Run the forward/backward elimination algorithm.
- Step 8: Run gradient descent on the final subset of modes. Make sure that this is done on the non-bandpassed data.
- Step 9: trim the data to remove edge artifacts.
- Step 10: use a least squares algorithm to find the estimate of **F**.

## III. RESULTS

We tested NS-DMD on simulations, electrophysiological brain data from multichannel recordings of local field potentials in the macaque brain, and from sea surface temperature (SST) data. NS-DMD recovers the underlying dynamics in the simulations. In the electrophysiological brain data, we show a rich set of modes that are active during different periods of performance of a cognitive task. In the SST data, NS-DMD recovers seasonal modes along with El Niño modes. Hyperparameters are described for all applications in Appendix B. Further simulations, which require optional steps of NS-DMD or are of interest to specific communities, are included in the supplementary material (Sec. S2).

### A. SIMULATIONS

Synthetic data is generated from the following generic equation:

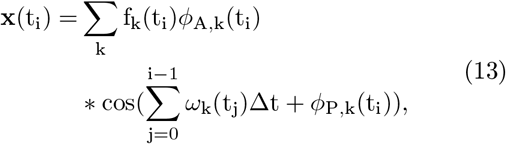

where **x**(t) is the data consisting of a vector of channels, f_k_(t) is the amplitude modulation of all channels, *ω*(t) is the time varying angular frequency, *ϕ*_A,k_ is the time varying normalized amplitude (|*ϕ*|≡ 1), *ϕ*_P,k_ is the time varying phase, and Δt 0.001 is the time delay between snapshots. This is the more explicit form of the simplified form of the data in Eq. 4, which can be seen by expressing the cosine in terms of exponentials and combining with *ϕ*_A,k_ to form **S**.

#### 1) Non-Stationarity in Multiple Assemblies

The simplest case of interest is when multiple assemblies switch on/off with non-constant amplitudes, corresponding to the following simplification of Eq. 13:

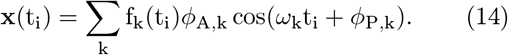

We construct a group of modes that are active in the beginning, a group of modes that are active towards the end, a switching period where multiple modes coexist, and a mode with non-constant amplitude. The assemblies, labeled from A to D, construct the data shown in Fig. 3A. Assemblies A and B operate with a constant amplitude for 1700 ms, assembly C turns on with a constant amplitude at t =1300 ms, and assembly D’s amplitude follows the shape of a Gaussian distribution starting at t =1300 ms. All assemblies initiate and decay with a fixed timescale. The exact shape of **F** is shown in Fig. 3D.

**FIGURE 3:**
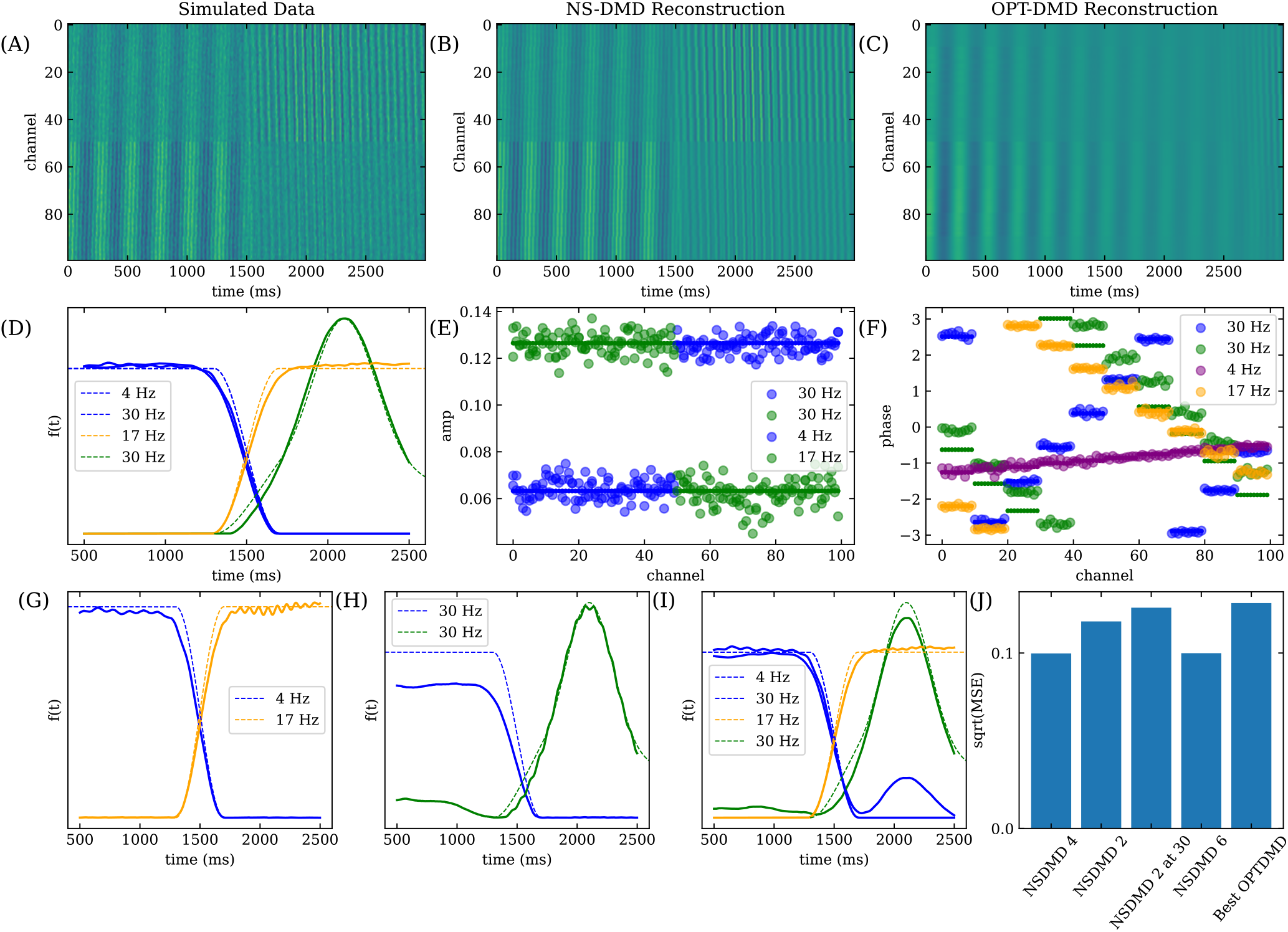
Multiple assembly simulation. The simulated data, generated from 4 assemblies A-D, is shown in (A). The frequencies are 4 Hz, 30 Hz, 17 Hz, and 30 Hz respectively. The dotted lines in (D) show the modulation f(t) for each assembly, and the small solid circles show the amplitude and phase of **Φ** in (E) and (F). Gaussian noise with a standard deviation of 0.1 is added independently to each channel and snapshot. (B) shows the reconstruction of the data with NS-DMD with 4 modes per window. (C) shows the reconstruction with OPT-DMD with a rank of 4, which had the smallest reconstruction error out of ranks from 2 to 10. (D) compares the true (dotted lines) and fit **F** (solid lines) for each mode/assembly. (E) and (F) compares the true (small, solid dots) and fitted (large, transparent dots) **Φ** amplitudes and phases for each channel. From this, we see the accuracy of the underlying modes, f(t), and reconstruction of the data when using NS-DMD with an optimal number of 4 modes per window. The inaccuracy in the green mode in (F) occurs since the phase is compared at t = 0 ms; the phase is most accurate at t ≈ 2000 ms when the mode is active, and any small inaccuracy in frequency will grow when analyzing the phase at a large time difference away. We compare the the true (dotted lines) and fit **F** (solid lines) when running NS-DMD with 2 (G), 2 and an initial guess of ±30 Hz (H), and 6 (I) modes per window. In general, we find that 2 modes per window fits lower frequency modes well. Forcing the initial guess to higher frequency modes, as in (H), leads to higher deviations from the ground truth. Using 6 modes per window, as in (I), leads to overfitting at 2000 ms, which may be expected due to the similarity of the constructed ±30 Hz modes. The reconstructed error for each number of modes per window is shown in (J), where the error is on the order of the Gaussian noise. The error is worse when using less number of modes due to an underfitting of the data. The best performance with OPT-DMD is still worse than underfitting NS-DMD models. Surprisingly, the error is still fairly small considering the blurred reconstruction in (C).

The frequency *ω*_k_*/*(2*π*) is 4 Hz, 30 Hz, 17 Hz, and 30 Hz for modes A-D respectively. Assemblies B and D have the same frequency, but with different spatial distributions. For A and B, the spatial amplitude is 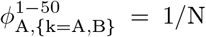 and 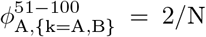, where the superscript labels the channel number and N is a normalization such that |*ϕ*_A,k_| = 1. The amplitudes of assemblies C and D are reversed such that 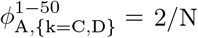 and 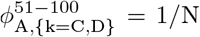. The exact amplitudes are shown in Fig. 3E. The spatial phases *ϕ*_P,k_ for every assembly form ten groups of ten channels. The phases are constant within each group of channels. Assembly A has a temporal delay *ϕ*_P,k_*/ω*_k_ from 0 ms to 30 ms across all groups; Assembly B has a delay from -20 ms to 30 ms; Assembly C from 50 ms to 0 ms; and Assembly D from 30 ms to -10 ms. The phases are shown in Fig. 3F. Each channel receives independent white noise with a standard deviation of 0.1.

We run NS-DMD with 4 modes per window, and the reconstruction of the data is shown in Fig. 3B. The reconstruction error is 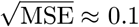 (Fig. 3J), the same as the added noise. Comparing the true and fitted **f** in Fig. 3D, NS-DMD recovers the correct mode amplitudes, including the non-stationary amplitude in assembly D. The spatial amplitudes and phases in Figs. 3E and 3F show recovery of the correct underlying modes. Overall, this simulation shows that NS-DMD can capture the underlying modes and temporal variations of a nonstationary linear dynamical system.

In practice one does not know the optimal number of modes per window. If instead we run NS-DMD with a smaller number of modes than optimal, 2 per window, it will only be able to capture half of the modes at any timestep. Due to the switching of assemblies, we expect NS-DMD to recover one early assembly (A or B) and one late assembly (C or D). NS-DMD recovers the 4 Hz and 17 Hz mode, as shown in Fig. 3G. This occurs due to the procedure; in the first window after the 4 Hz mode “turns off,” the initial guess with OPT-DMD is still 4 Hz. OPT-DMD then converges to the closest solution, which happens to be 17 Hz. The reconstruction error (Fig. 3J), is marginally worse, as expected since only half of the modes are captured.

If the initial guess for the first window is forced to be 30 Hz, then NS-DMD finds the two 30 Hz modes (Fig. 3H)). NS-DMD performs noticeably worse when finding **F**, but it still finds the correct trend. The low amplitude bias of the 30 Hz mode occurs due to a lack of an estimated 4 Hz mode, which can bias **X** to more positive or negative values for short (< 50 ms) windows. We verify this by rerunning this model with only assemblies B and D, where **F** is found correctly. The reconstruction error in Fig. 3J indicates a preference for lower frequency modes.

If we run NS-DMD with 6 modes per window, NS-DMD recovers 4 modes that reconstruct the data. The fitted and recovered amplitudes **F** in Fig. 3I show that NS-DMD performs well. Around t=2000 ms, both 30 Hz modes have a non-zero f(t) which occurs due to their similar construction. The reconstruction error is on par with the amount of noise (Fig. 3J).

Lastly, we run OPT-DMD with ranks ranging from 2 to 10 and find the best performance with a rank of 4. The reconstruction is shown in Fig. 3C, where the original structure in the simulated data is blurred. Despite this, the reconstruction error is fairly small (Fig. 3J). However, the reconstruction error is worse than all NS-DMD models.

#### 2) Smoothly Varying Time Dependent Modes

We now consider the case of smoothly varying time dependent modes (Fig. 4A). Allowing a time-varying frequency corresponds to the following simplification of Eq. 13:

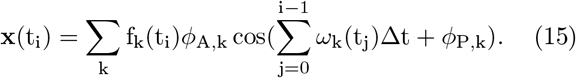

We simulate three assemblies whose frequencies vary linearly in time around 17 Hz, 27 Hz and 33 Hz, where the true frequency *ω*_k_*/*(2*π*) is shown in Fig. 4C. The 27 Hz assembly occurs early in time while the 33 Hz assembly occurs late. The 27 Hz assembly appears with a linearly varying amplitude for the duration of the dataset. The exact amplitudes **f** (t) are shown in Fig. 4B.

**FIGURE 4:**
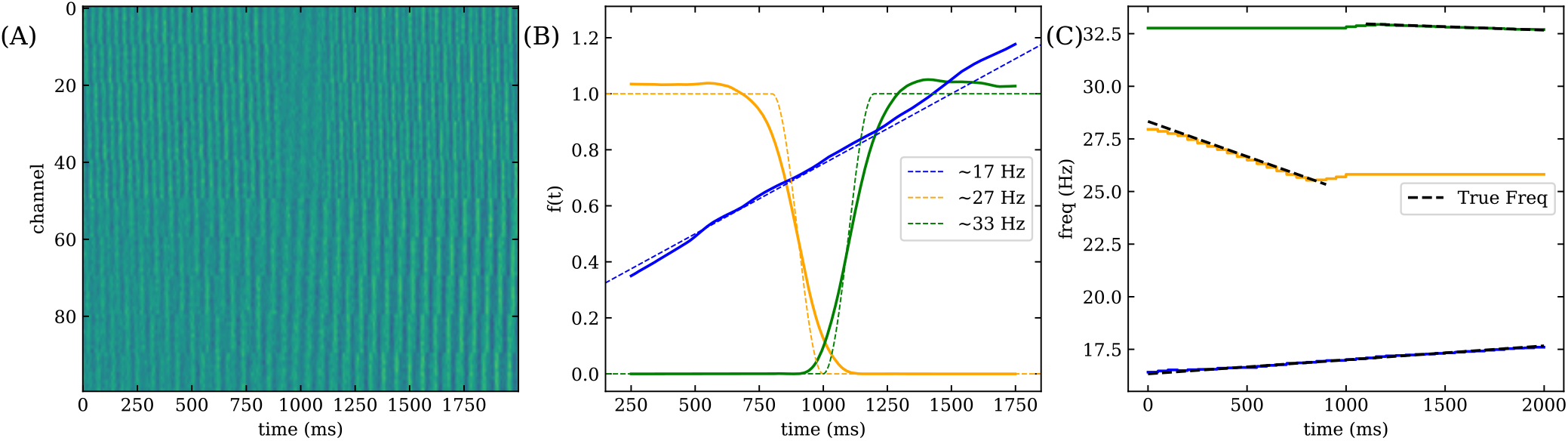
(A) Synthetic data for the smoothly varying time dependent mode simulation. (B) A comparison of the true (dotted lines) and fitted **F** (solid lines) for three assemblies with time dependent frequencies. NS-DMD recovers the true modulation trends. (C) Comparison of the true frequencies (dotted lines) with the fitted frequencies (solid lines) for each drifting assembly. Note that when **F** = 0 for a given mode in (A), the frequency is undefined. While not shown, we again compare the performance of OPT-DMD to NS-DMD. The reconstruction error is 0.1 for NS-DMD and ∼ 0.12 for OPT-DMD. Again, the visual features are blurred compared to a visually accurate reconstruction from NS-DMD.

NS-DMD recovers the correct **F** (Fig. 4A), and time dependent frequency (Fig. 4B). Note that the frequency is undefined when f(t) = 0, indicated in Fig. 4B by ending the dashed lines.

We next allow for the spatial amplitude to fluctuate instead, corresponding to the following simplification of Eq. 13:

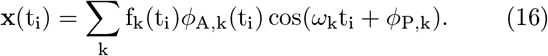

The frequencies of two assemblies are fixed to 17 Hz and 30 Hz. The spatial amplitude is shown in Figs. 5A and 5D respectively, where channels 1-50 and 51-100 are grouped together with the same change in *ϕ*_A,k_ over time.

**FIGURE 5:**
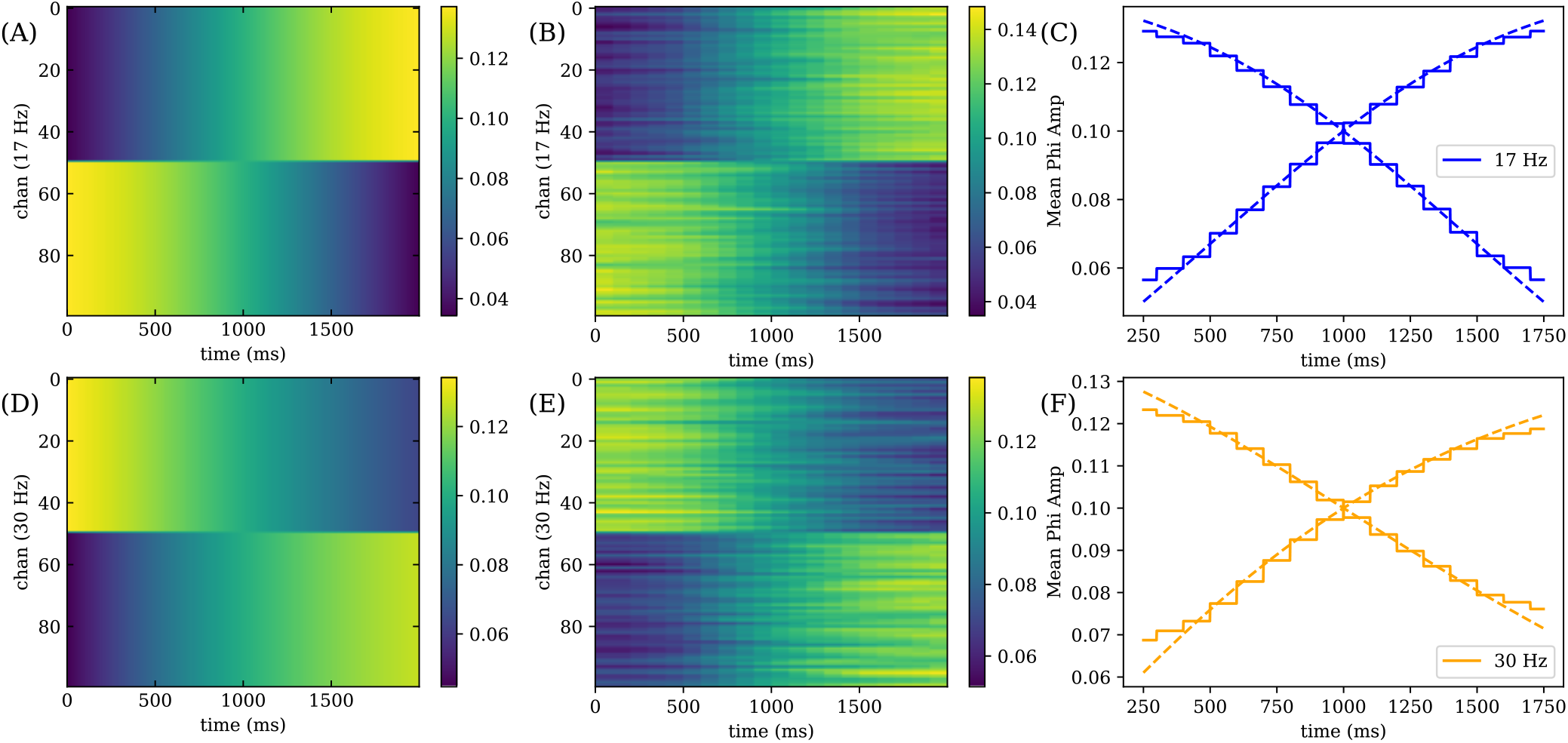
(A) and (D) True spatial amplitude distribution for time dependent spatial modes at 17 Hz and 30 Hz, where channels 1-50 and 51-100 are grouped and vary the same way. (B) and (E) Fitted spatial amplitude distribution for the 17 Hz and 30 Hz assemblies. (C) and (F) Comparison of the true (dotted) and fitted (solid) mean *ϕ* amplitude for channels 1-50 and 51-100. We find that NS-DMD recovers the correct spatial amplitudes *ϕ*_A,k_, where the staircase pattern occurs due to the 100 ms stride of NS-DMD. When running OPT-DMD, the reconstruction error is ∼ 0.12 compared to 0.1 for NS-DMD. The visual features are blurred compared to NS-DMD.

NS-DMD recovers the correct spatial amplitudes, shown in Figs. 5B and 5E with a step like variation due to discrete windows. To aid visualization, we plot the average amplitude for each mode and group of channels, 1-50 and 51-100 in Figs. 5C and 5F respectively.

We run OPT-DMD as well for both simulations of fluctuating frequency and spatial modes. The reconstruction error is worse for OPT-DMD at about 0.12 compared to 0.1 for NS-DMD. Like in the example in Fig. 3, the visual features are blurred with OPT-DMD (not shown), indicating the need for NS-DMD to recover the correct underlying modes.

### B. APPLICATION TO ELECTROPHYSICAL BRAIN DATA

We apply NS-DMD to local field potentials (LFP) [7], an invasive measurement technique in which electrodes measure electric potentials deep inside the brain. In general, the LFP power decays approximately as a power law, with an exponent between −1 and −2 [24]. A wealth of literature has found correlations between behavioral parameters and time-varying power in various frequency bands of the LFP [21, 54, 27]. Other research suggests the possibility of cross-frequency coupling [14, 64, 11, 38] as a top-down mechanism of control.

The LFP is typically analyzed using standard timeseries procedures, such as Hilbert or spectrogram analysis [10, 13], coherence analysis [58], and Granger Causality [55, 52]. While these methods are useful for understanding the structure of the data, they do not lead to a dynamical systems model of the brain. Others have argued for the use of a Koopman operator approach [37]. A DMD approach has been applied to sleep activity [4], revealing sleep spindle networks. Given its non-stationary spectral properties and the potential for the application of DMD in analysis of brain activity, LFP data are a perfect candidate for NS-DMD.

We demonstrate that NS-DMD can find consistent, repeatable spatial modes that co-activate intermittently in correspondence with the task. Modes activate and deactivate in correspondence with task events, and they cluster in different areas of the brain. Further, some clusters show consistent phase differences between brain areas, indicating information flow. NS-DMD is also able to recover results from standard time-series analysis.

#### 1) Dataset

We apply our methods to LFP data collected in the Buffalo lab from a macaque monkey performing a variant of the Wisconsin Card Sorting Test [19]. Out of 4 non-human primates performing the task, two have electrodes (FHC and Alpha Omega) implanted for neural recordings. A single subject is chosen for analysis and comparisons between methods. The subject is an adult female rhesus (Macaca Mulatta), aged 9 with a weight of 9.1 kg. The subject was headfixed with a titanium rod in a dimly lit room. The subject was positioned 60 cm away from a 19-inch CRT monitor, with 33 degrees by 25 degrees of visual angle. Stimuli were presented on the screen with software (NIMH Cortex). All procedures were carried out in accordance with the National Institutes of Health guidelines and were approved by the University of Washington Institutional Animal Care and Use Committee.

Trials are initialized when the animal fixates on a cross in the middle of the screen. The monkey must then choose one correct image out of four simultaneously presented images based on an uncued rule. The rule is discovered by trial and error. Each image has one of four possible shapes, colors, and patterns, and the rewarded rule is one of the 12 possible visual features. Animals are rewarded with a juice/chow mixture for 1400 ms if correct, and given a 5000 ms timeout period if incorrect. The rule remains fixed across consecutive trials while the monkey learns it; after 8/8 or 16/20 correct trials, the rule spontaneously changes.

The LFP is recorded for ∼ 3 hours per session across several months using 220 electrodes implanted in multiple locations throughout the brain, including hippocampus and prefrontal cortex; we focus here on data from a single day. We ignore 17 electrodes that are dominated by noise, determined based on an unusually large amount of 60 Hz wall noise. We neglect trials where the remaining electrodes experience random artifactual bursts, or LFP activity that reaches the maximum or minimum possible recording limit of the electrode. This leaves us with 896 trials recorded on 203 electrodes to analyze. NS-DMD is applied to the raw data, normalized by z-scoring each electrode’s signal in the 1-40 Hz range independently for each trial.

#### 2) NS-DMD on a Single Trial

We fit NS-DMD on a single trial of the LFP and analyze the resulting modes. There are non-zero modes during every section of the task. The amplitude f(t) of up to 10 modes within the 2-7, 12-17, 22-27, and 32-37 Hz modes are plotted in Fig. 6(A). Other modes exist, but are not shown for concision. Some modes span very long stretches of the trial, while others turn on or off relative to task events. The red mode of the top plot and the purple mode of the bottom plot of Fig. 6(A) are selected for further analysis. In Fig. 6(B) and Fig. 6(C) the spatial amplitudes |*ϕ*| and phases ∠*ϕ* are shown for the red mode (top) and purple mode (bottom). The spatial amplitude |*ϕ*| is plotted. The spatial modes showcase patterns across the brain. In Fig. 6(C), the phases show significant differences across various areas of the brain, indicating that some areas lead or lag behind other areas in these particular modes.

**FIGURE 6:**
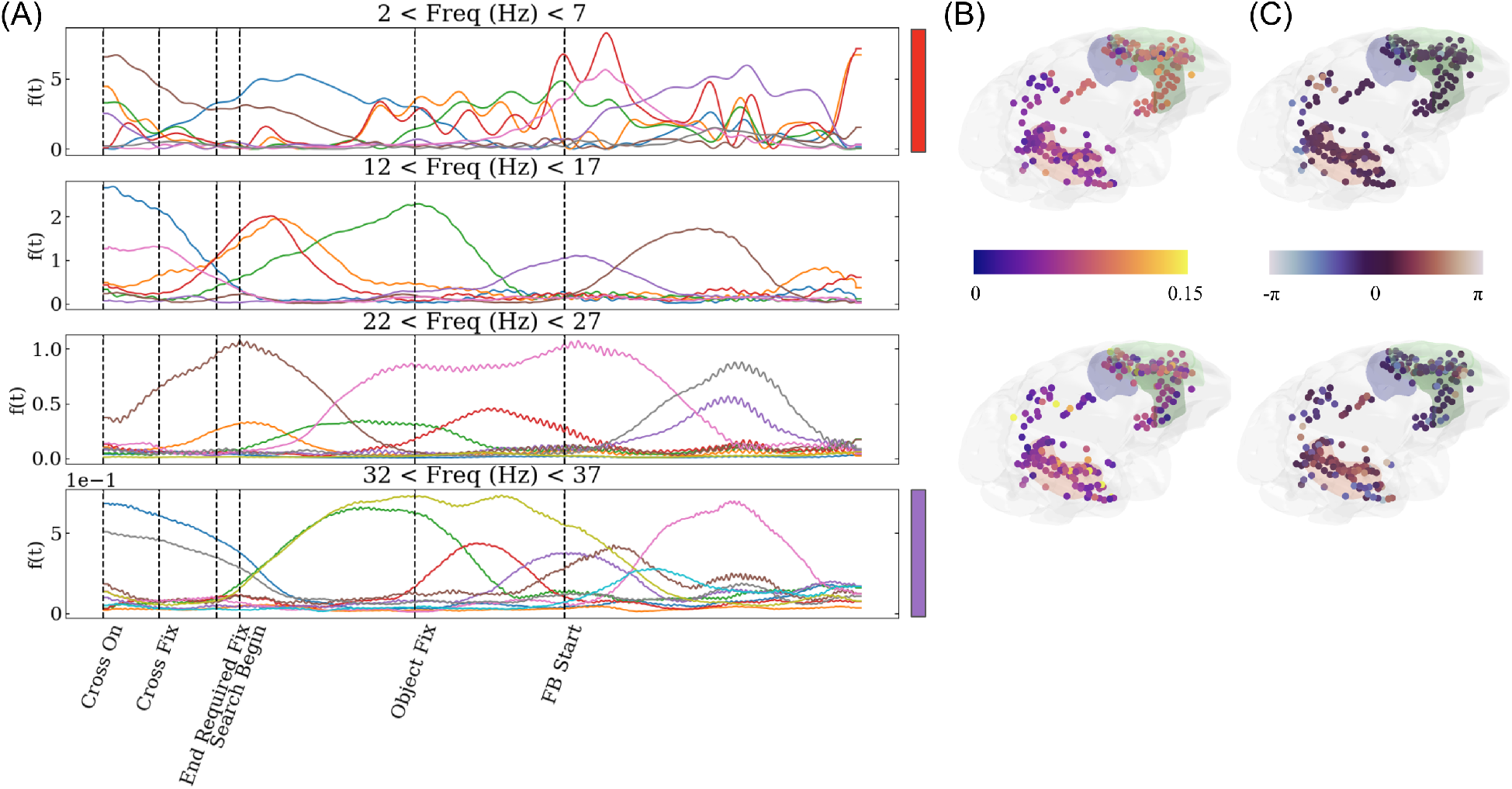
Example NS-DMD modes for the first trial of the LFP data. (A) shows the mode amplitudes f(t), separated by frequency ranges: 2-7, 12-17, 22-27, and 32-37 Hz. Some modes persist for long periods of time, such as the red mode in the top two plots. Other modes persist for short periods of time, emphasizing task intervals, such as the red modes in the bottom two plots. (B) shows the absolute value of the relevant spatial amplitudes |*ϕ*| for the red (top) and purple (bottom) modes. There are large spatial spreads for each mode. (C) shows the relevant spatial phases ∠*ϕ* for the red (top) and purple (bottom) modes on a circular color scale. There are clear divides between the phases of different brain areas, indicating that some areas lead or lag behind others. This is especially true in the purple mode (bottom), where there is a phase difference between the red and green shaded regions. Shaded regions emphasize brain regions important for decision making and memory: hippocampus (red) and prefrontal cortex (green).

#### 3) Common NS-DMD Modes in All Trials

We then apply NS-DMD on all trials for 1500 ms after feedback begins, focusing on the differences between correct and incorrect trials in the 19-21 Hz frequency band. We perform k-means clustering [36] on the mean spatial amplitude |*ϕ*|, where each “point” in the k-means algorithm is the average |*ϕ*| when f(t) > mean(f(t)). We choose 3 clusters for both “correct” and “incorrect” modes. We further separate the data by performing k-means clustering with 3 modes using the temporal amplitudes **F** for each previously found group. This separates the modes into clusters that have distinct spatial distributions and temporal distributions. We select three groups out of the nine for both correct and incorrect trials for plotting.

The average spatial amplitude for each cluster is shown in Figs. 7A and 7B. Different clusters are associated with activity in different channels; some clusters have large amplitudes in single channels. The average global modulation 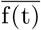 of each “correct”/”incorrect” cluster is shown in Figs. 7C (correct) and 7D (incorrect). In the correct trials, **F** separates into early and late modes. In the incorrect trials, there is a cluster with a large amplitude 1s after feedback is given. The overall amplitude of the “incorrect” modes is much larger than the “correct” modes.

**FIGURE 7:**
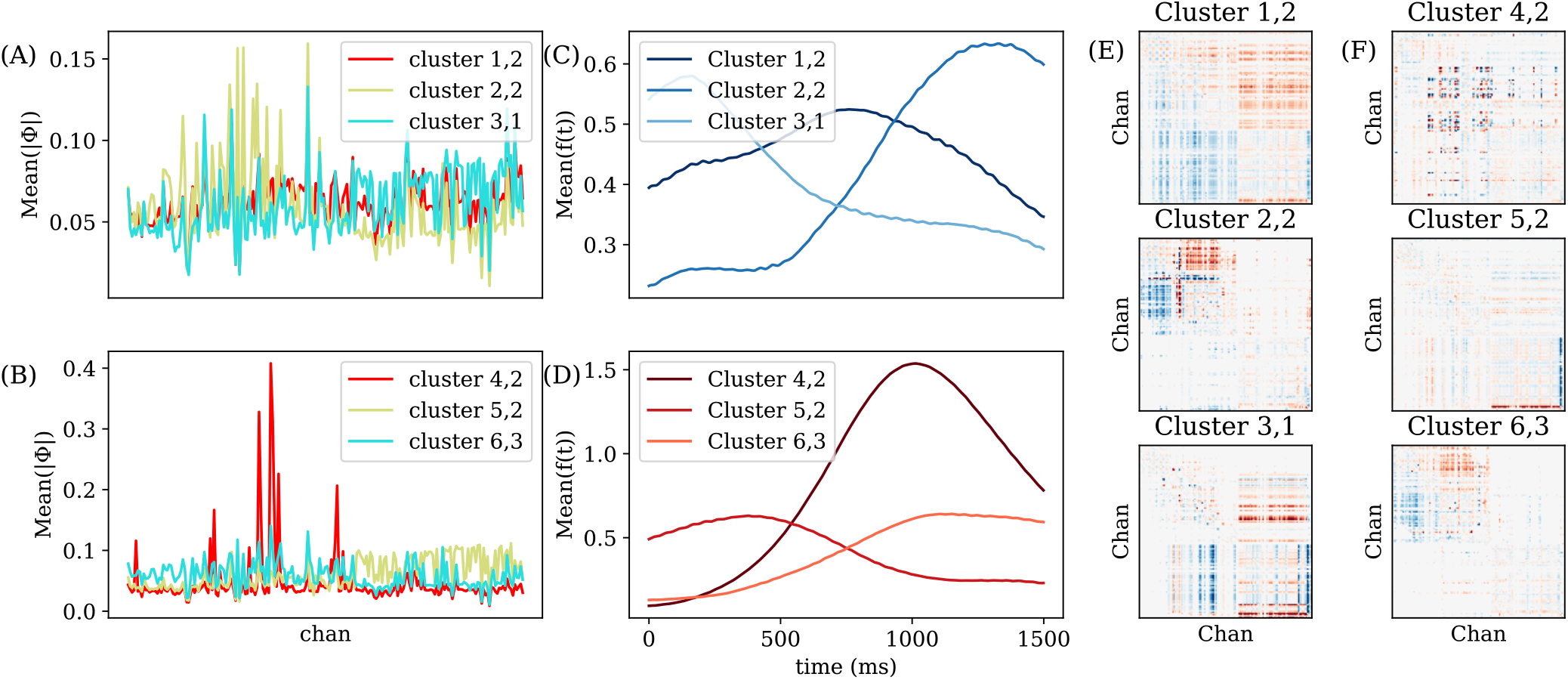
We group all modes corresponding to correct or incorrect trials into 3 clusters with K-means on the spatial amplitude (first index in labels). We further group each cluster into 3 additional clusters (second index in labels) based on f(t) for a total of 9 groups per correct and incorrect trials. 3 groups are selected for plotting, emphasizing consistent information flow during different periods after feedback. (A) and (B) show the average spatial amplitude of each cluster for the correct and incorrect clusters. (C) and (D) show the mean global modulations 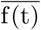 for each correct/incorrect cluster. Note that 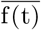 is averaged over many modes with independent f(t)’s. Thus, local maximums indicate when some, but not necessarily all, modes are large. (E) and (F) show the average phase difference between every pair of electrodes for correct/incorrect clusters. We average with Eq. 17, and the phase difference is set to 0 for every channel pair where the average phase amplitude is less than 0.4, corresponding to a p-value less than 3×10^−6^. The color limits are from −π/4 to π/4. The largest amount of widespread information flow occurs in the correct clusters between different brain areas, particularly directly after feedback. There is local flow in cluster 4,2. Interestingly, there is a similar cluster occurring in both correct (cluster 2,2) and incorrect (cluster 6,3) trials, where there is similar spatial, temporal, and phase difference activity.

We are interested in the phase difference between each channel pair, since this is indicative of information flow. The phase difference is averaged separately across correct and incorrect trials when 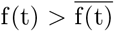. We represent the phase difference *θ* as a complex vector on the unit circle and average:

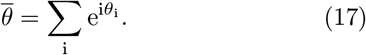

The amplitude of the average phase difference vector indicates how consistently the two channels are related. We focus on pairs of electrodes with large amplitudes where 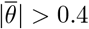. Significance is calculated from the work of [20]. For 76 vectors, which is the minimum number of vectors averaged in Fig. 7, a threshold of 0.4 corresponds to a p-value of 3 × 10^−6^. The average thresholded phase differences are shown in Figs. 7E and 7F. The “correct” clusters have the largest phase differences, suggestive of information flow between two groups of channels after correct feedback is given. Cluster 4,2 has sparse consistent phase differences, which occur during incorrect trials one second after feedback. There are two clusters, (2,2) and (6,3), which have very similar average patterns: they are similar across space and time, and the phase differences are shared. This shows that a common pattern emerges for both correct and incorrect trials.

#### 4) Comparison to Standard Analyses

To demonstrate the ability for NS-DMD to recover results from simpler methods, we compute analysis based on the Power Spectral Density, Hilbert analysis, and coherence.

First, for each electrode, we calculate the Power Spectral Density (PSD) to find the standard power law decay [7] using the Welch algorithm [62]. This results in the standard frequency power law in Fig. 8A. For NS-DMD, we average f(t) across all trials and times within overlapping 3 Hz frequency bands (Fig. 8B). The power law is recovered with a similar slope.

**FIGURE 8:**
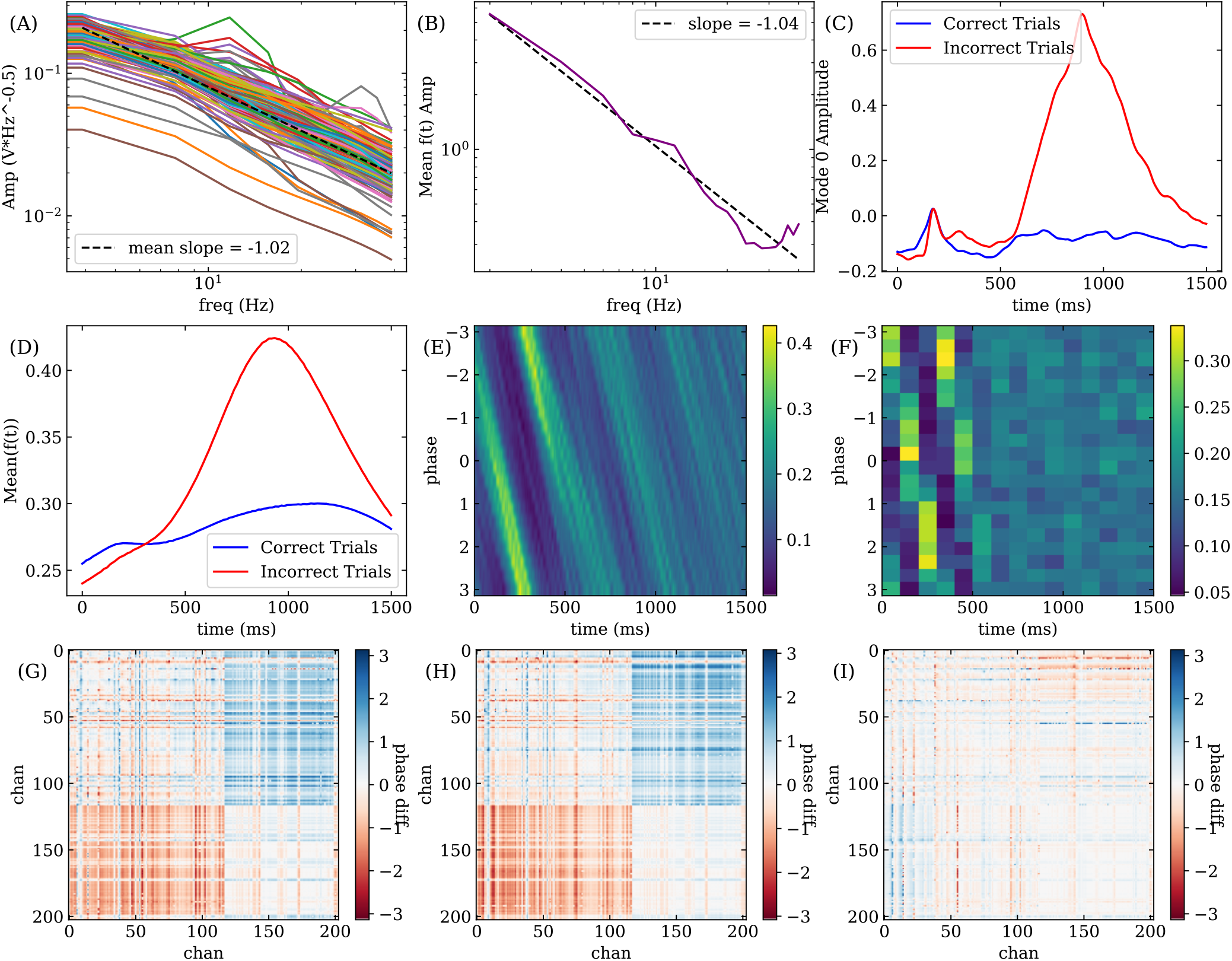
Comparisons of NS-DMD results with traditional analyses. (A) shows the Power Spectral Density plots for every electrode along with their mean power law decay. For the NS-DMD modes, (B) shows the average f(t) for every 3 Hz band along with the slope. NS-DMD finds a similar, but mildly different, power law decay to (A). After bandpassing and Hilbert transforming the data between 27 and 37 Hz, (C) shows the first PCA mode after feedback begins at t = 0 ms. For NS-DMD, (D) shows the mean f(t) for all modes between 27 and 37 Hz, averaged across correct and incorrect trials, matching the trend in (C). The difference in scale is due to the normalization in the Hilbert method. (E) shows the normalized histogram of phases, determined by the Hilbert transform between 2-4 Hz, across all trials for each time point. The normalized histogram of the NS-DMD phases between 2-4 Hz is computed across all trials for each time point in (F). To compare with standard coherence, we take the average phase difference between every pair of channels for incorrect trials in (G). Coherence is calculated at 3.5 Hz for a window of 0-500 ms between every pair of channels, and the phase difference is averaged with Eq. 17. All average phase values are set to 0 when the amplitude of the average phase vector is less than 0.1. (H) shows the average phase difference between every pair of channels for incorrect trials, computed with NS-DMD. Modes are averaged, where the modes are between 2-4 Hz and where 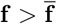 within 100-300 ms. NS-DMD finds a similar answer to (G) with a mean squared error of 0.17. The difference is shown in (I).

Second, we compare NS-DMD to a traditional Hilbert analysis, where one typically analyzes a frequency band of interest. We bandpass the data to 27-37 Hz, Hilbert transform every channel, and take the absolute value of the resulting signal.We concatenate the trial and time dimensions, and run Principal Component Analysis (PCA) to reduce the dimensionality. By averaging the projection of the data onto the first mode for all correct/rewarded trials and all incorrect/unrewarded trials, we find separate phenomena for each trial type (Fig. 8C).

For NS-DMD, we average f(t) across correct and incorrect trials for all modes that occur within 27 and 37 Hz. We find the same amplitude trends in Fig. 8D, where a large amplitude occurs in incorrect trials one second after feedback. The scale is different in Figs. 8C and 8D due to normalization. The similarity between overall trends indicates that NS-DMD recovers similar results to standard Hilbert analyses.

Next, we compare NS-DMD to the phase of the Hilbert transform. Jutras et al. (2013) [26] showed that eye movements are associated with phase resets. We explore whether feedback events also cause a phase reset. We bandpass and Hilbert transform every channel between 2 and 4 Hz. The angle of the Hilbert transform finds the instantaneous phase for every millisecond of every trial. We find a consistent phase shortly after feedback begins. A phase histogram for an example channel is shown in Fig. 8E. We compare the phase in the NS-DMD modes, ∠*ϕ*, for the same example channel. The histogram of phases of all modes between 2 and 4 Hz is shown in Fig. 8F. The same phase reset can be seen in the NS-DMD phases.

Lastly, we compare NS-DMD to an example coherence analysis. We calculate the coherence between every pair of electrodes with the Welch method at 3.5 Hz for 500 ms after feedback begins. We then find the average phase difference between all pairs of channels across correct and incorrect trials, given by Eq. 17. We consider only phases for which the magnitude of the average phase vector 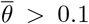. The resulting spatial phase difference map for incorrect trials is shown in Fig. 8G. For NS-DMD. We find the phase difference between all pairs of electrodes for modes between 2 and 4 Hz and when 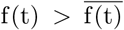 between 100 and 300 ms. We average the phase differences with Eq. 17. The resulting spatial map in Fig. 8H matches the coherence analysis. The difference is shown in Fig. 8I, highlighting some small, local differences. The mean squared error (MSE) is 0.17.

### C. APPLICATION TO SEA SURFACE TEMPERATURE

We apply NS-DMD to sea surface temperature (SST) data, where known global frequencies exist. SST data (NOAA Optimum Interpolation (OI) Sea Surface Temperature (SST) V2 [48]) is collected via satellite, and it is averaged weekly from 1990 to 2016 on a 180 by 360 grid across all longitudes and latitudes. After flattening and removing land locations, we end with a length 44219 vector for 1455 weeks. The data for each recording location are normalized by z-scoring across weeks.

The power spectrum density (PSD) of a sample location in the Pacific Ocean is shown in Fig. 9(A), where there appears to be different frequencies: there is a high amount of power once per year, a smaller amount of power twice per year, and a fluctuating amount of small power less than once per year.

**FIGURE 9:**
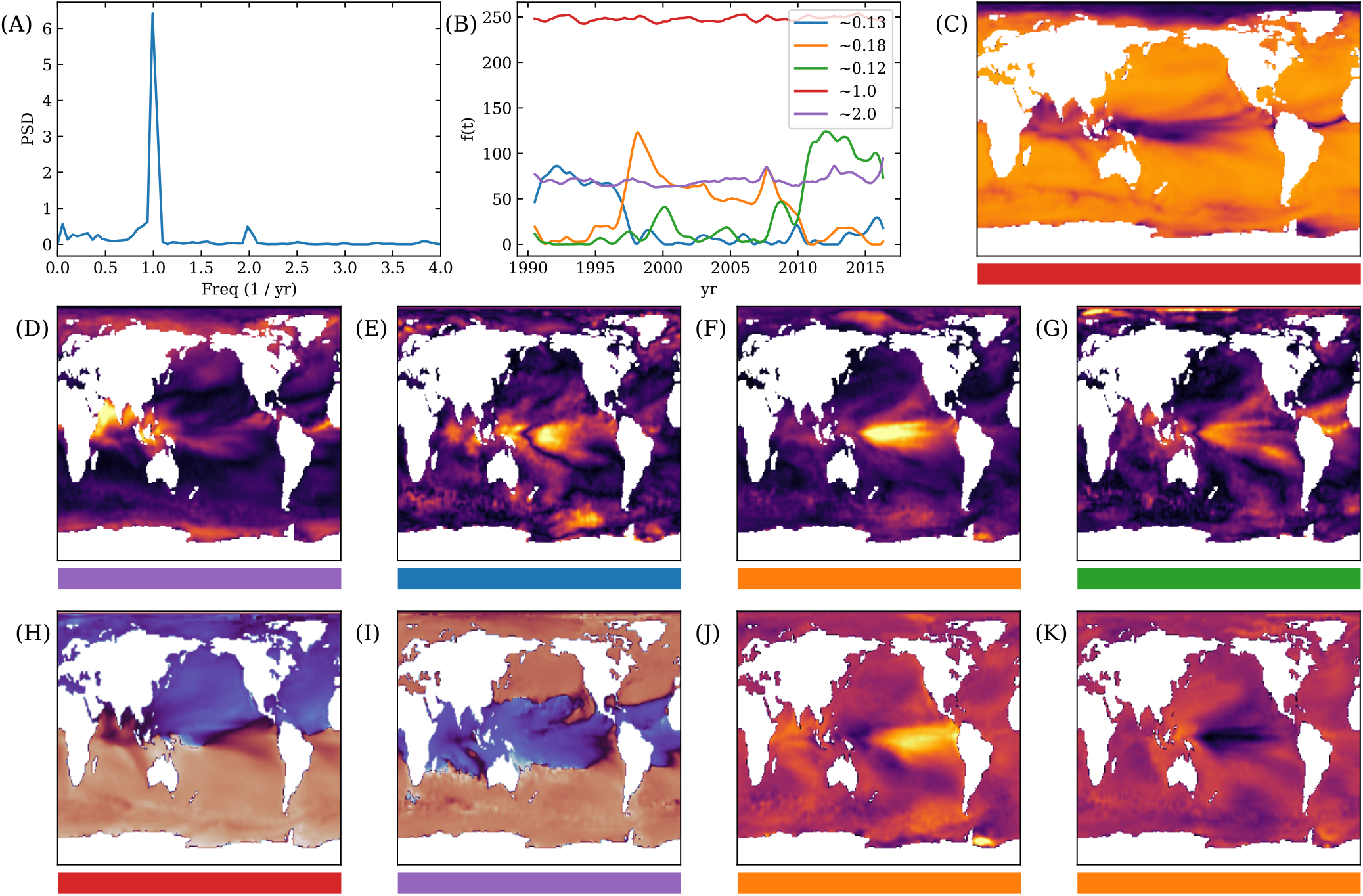
Application of NS-DMD on Sea Surface Temperature data ranging from 1990 to 2016. (A) shows the power spectrum density for a random location in the Pacific Ocean. (B) shows the amplitudes of the 5 resulting NS-DMD modes **F**. The legend highlights the average frequencies for each mode in units of cycles per year. Note the two stationary modes that occur due to seasonal effects. The other three modes tile the duration of the dataset and are most likely El Niño and La Niña modes. For (C) through (K), the horizontal bar showcases which mode it comes from, in terms of the colors in (B). (C) shows the spatial amplitude |*ϕ*| for the once a year mode. The relative phase ∠*ϕ* is shown in (H), where the Northern and Southern hemispheres are a *π* phase apart. The spatial amplitude for the twice a year mode is shown in (D) and the relative phase is shown in (I). This mode appears to be strongest in equilateral latitudes, particularly in the Indian ocean. The phase appears to be *π* offset between latitudes near the equator and outside the equator. The boundaries appear consistent with the tropic of Cancer and Capricorn, where the sun may be directly above the Earth’s surface. (E) through (G) show the spatial amplitudes for the El Niño and La Niña modes, where the strength is particularly strong in the Pacific ocean. The data is reconstructed from solely the orange, 0.18 times a year mode, and the resulting reconstruction for a week in January in 1998 and 1999 are shown in (J) and (K) respectively. The ocean temperatures are known to have exhibited a very strong El Niño event in 1998 and a very strong La Niña event in 1999.

From the PSD, we estimate that 6 modes per window are sufficient to reconstruct the data. We run NS-DMD with 6 modes per window, a window size of 150 weeks, a stride of 25 weeks, and we guess the 6 modes have frequencies of ±1, ±2, and ±0.12 per year. We compute for the first 1400 weeks. Specific parameters are labeled in App. B. We find that 5 modes, that span the entire 1400 weeks, reconstructs the data well. The cosine distance between the reconstruction and original, z-scored data is about 0.92.

The amplitudes **F** are shown in Fig. 9(B) for the 5 modes. There are two modes that exist with a constant amplitude at once and twice per year. These modes correspond to seasonal changes in the SST. The once a year mode is present across the entire ocean (Fig. 9(C)). Examining the phase in Fig. 9(H) shows a *π* offset in the phase from the Northern and Southern hemispheres, occurring due to the tilt of the Earth. The twice a year mode occurs in equilateral locations with an emphasise in the Indian Ocean (Fig. 9(D)). This also occurs due to the seasonal tilt of the Earth, where equilateral locations undergo a twice a year heating event when the Sun shines most directly on it. The phase of the twice a year mode is shown in Fig. 9(I), where we see a *π* offset between the equilateral and non-equilateral locations around the Tropics of Cancer and Capricorn.

The other three modes in Fig. 9(B) are El Niño and La Niña modes, and occur with frequencies of 0.12, 0.13, and 0.18 per year. El Niño and La Niña are typically referenced to occur about once every 6 years, or with a frequency of 0.167. These three modes span the 1400 weeks, where when one mode turns off, another turns on. The spatial amplitude is shown in Figs. 9(E)-(G), where one can see the characterizing large amplitude in the equilateral latitudes in the Pacific ocean. To confirm, we analyze the orange, 0.18 times a year mode in 1998 and 1999, which were known as exceptionally strong El Niño year and La Niña years. By looking at the orange mode during a week in January of 1998 and January of 1999, we see a relative increase and decrease in the temperature near the equator in the Pacific Ocean (Figs. 9)(J) and (K).

We highlight that NS-DMD is applicable to sea surface temperature data, and it can find modes of particular interest. It is appropriate for this problem due to the combined stationary and non-stationary modes. The seasonal changes in temperature are stationary, but the El Niño and La Niña effects are non-stationary.

Other groups have approached SST data with DMD like methods: [32] uses multi-resolution DMD, [22] uses Time-varying Autoregression with Low Rank Tensors (TVART), and [50] uses BOP-DMD to analyze SST data. They have all shown success in finding modes correlating with El Niño and La Niña in the Pacific Ocean. Our approach, however, allows us to find differences in the spatial amplitude during different years. The modes attributed with El Niño and La Niña turn on and off during two specific years, indicating that the structure may be changing slightly.

## IV. DISCUSSION

We introduce a novel method for analyzing time-series data: Non-Stationary Dynamic Mode Decomposition. NS-DMD builds on previous DMD methods by including global modulation and time dependent modes. Thus, any improvement to standard DMD algorithms can be easily integrated into NS-DMD. NS-DMD accurately discovers modes that well explain data across a range of simulated settings. III-A. Naturally, this method is best suited for data that includes low-rank spectral features. It is possible to run DMD on any time-series data, since any signal can be decomposed into a Fourier series basis set of sines and cosines.

NS-DMD can be useful in many empirical settings. This is because many systems elicit non-stationary behavior. For such systems, NS-DMD is better suited than previously proposed methods that assume stationary properties. In the present work we demonstrated this in data from large-scale neural recordings and from recordings of SST. As shown in Sec. III-B, for these empirical data, NS-DMD extends and subsumes standard methods, such as spectrograms, wavelet transforms, or coherence. However, while these methods work on individual recording locations or can be combined with global dimensionality reduction techniques, the main benefit of NS-DMD is to simultaneously gather spatial information, spectral information, and growth and decay of all modes.

NS-DMD can capture drifting components since it allows for modes to modulate slightly over time. It is then useful to combine modes into one single drifting mode instead of defining multiple new processes. Among its limitations is the fact that NS-DMD requires careful choice of the correct values of hyperparameters. If the similarity threshold is too tight, a single mode will be parsed into many chunks at different short intervals of time. If the similarity threshold is too broad, we lose the ability to distinguish between mode switching and time dependent modes. Expert knowledge can help with hyperparameter choice. Alternatively, to decide on hyperparameters, we recommend directly analyzing modes from consecutive windows. E.g., if the modes gradually change, the similarity hyperparameters should allow these modes to be defined as similar. Or if the modes turn on or off rapidly, a tighter similarity hyperparameter can be used.

### A. RELATED WORK

Many previous approaches exist to fit non-stationary systems that are closely related to NS-DMD. In this section, we describe some of this previous related work, and draw distinctions between these previous approaches and the NS-DMD approach developed in the present paper. These include hidden Markov models [15, 6, 59], and time-varying autoregressive models [23]. Generally, these methods contrast with our approach in assuming discrete state transitions and do not fully capture the dynamics in terms of identifying spatiotemporal modes of the system. Piece-wise Locally Stationary Oscillation models [57] and state-space multi-taper methods [28] focus on non-stationary estimates for univariate timeseries recordings. However, while these approaches can independently model individual components of a multivariate time-series, they cannot find low dimensional spatial modes combining components.

Variations of DMD [31] seek to address the full spatiotemporal dynamics, including finding reduced dimensionality spatial modes of oscillatory dynamics, across large, multi-variate systems [35, 32]. While powerful in principle, DMD is highly sensitive to noise [9, 2], thus generating biased and inaccurate estimates of the dynamics. Optimized DMD provides the most stable and biased estimate of a DMD model [2], with the bagging, optimized DMD (BOD-DMD) method [50] improving the method even further by providing uncertainty estimates of the DMD fit. But these DMD methods still fail when the generating dynamical system switches between different approximately linear regimes [41]. Within the Koopman framework, Macesic et al. [35] introduce two methods for dealing with rapid switches in the underlying system as well as continuously varying time-series, although expert knowledge of the system is needed to introduce observables that linearize the dynamical system. One variation of particular interest, Multi-Resolution DMD [32], specifically accounts for non-stationary time-frequency data. In this approach, one seeks DMD modes at different frequency scales, and with smaller and smaller windows, which increases the temporal resolution. The method successfully identifies non-stationarities, e.g., the El Niño effect in ocean temperature data. One downside, however, is that expert knowledge of the appropriate window size at different scales is required. This also assumes that lower frequency modes are more stable in time. While this may be appropriate for some systems, such as ocean temperature, this is not generally true for all non-stationary systems.

Switching Linear Dynamical Systems (SLDS) [16, 12, 34, 8], assumes a Markov process that switches between discrete, linear systems. A recurrent version was developed for neuroscience applications in [17]. These types of models find discrete states and transitions between them. NS-DMD can improve on this by allowing the states themselves to modulate over time, i.e. with a continuously variable amplitude or frequency. It also allows for independent transitions in individual modes or spatiotemporal components of the dynamics, without requiring an entire state to transition.

Another recent addition to the toolbox of methods for time varying systems is Latent Factor Analysis via Dynamical Systems (LFADS) [60, 42], in which smooth, low-dimensional dynamics are inferred using deep learning based on initial conditions and inferred inputs. While LFADS was first developed for neuronal spike counts, modeled as point processes driven by underlying latent dynamics, recent work [40] has extended LFADS to continuously varying signals. This method does not assume linear dynamics and instead uses recurrent neural networks to find low dimensional factors. Since NS-DMD uncovers linear approximations, some representations may be easier to interpret, such as the leading and lagging of individual spatial areas.

Lastly, a recent approach to non-stationary time-frequency data is Time-Varying Autoregression with Low-Rank Tensors [22], which successfully identified low rank modes and crossover points for constantly evolving data. This method is extremely similar to NS-DMD, even finding global modulations of individual modes. However, instead of finding dynamics in the form of Eq. 2, they find the global modulation of two spatial components; there is a lack of frequency and phase of each spatial mode. E.g., if one is interested in the global modulations of modes at a particular frequency or if one is interested in the phases of the spatial distribution, NS-DMD can be more informative.

### B. FUTURE DIRECTIONS

There are a number of potential additions for improving the effectiveness of NS-DMD. First, we implemented Optimized Dynamic Mode Decomposition [2], which is known to be more robust to noise than the standard DMD method. In the future, we plan on adding Bagging Optimized DMD (BOP-DMD) [50] to NS-DMD. In BOP-DMD, one runs OPT-DMD many times for each window to find statistics of each mode. BOP-DMD also can provide a metric to quantify uncertainty as it automatically produces probability density estimates for the modes, eigenvalues and loadings of the DMD approximation. This could aid with some cases where excess, poorly estimated modes are removed.

In the gradient descent method, we have imposed non-negativity and continuity by manually setting negative values to zero and smoothing across time. Given advances in Non-Negative Matrix Factorization (e.g. [33]), we believe it possible to further optimize the gradient descent method to implicitly restrict the values. We also believe we can implicitly add averaging into the methods.

Another assumption of NS-DMD is that the data are real. If the data are instead complex, one can easily transform it by squaring the magnitude. In the future, however, we plan on generalizing the gradient descent method to allow for complex inputs.

In allowing modes to disappear for extended periods of time, one expects in general that they can reappear with a phase unrelated to the previous appearance. However, NS-DMD retains phase information about modes. As we explore in the supplementary simulations (Sec. S2-C), under these conditions NS-DMD will either add an entirely new mode or mix modes. Ideally, phase should reset anytime that **F** reaches 0. In the future, we plan on implementing this by checking similarity in non-consecutive windows, allowing for merging of modes differing only by phase when delayed by large time intervals. This should remove any mode mixing and help with interpretability.

Lastly, we have implemented relatively simple feature selection algorithms. Given the large body of work in this area, we plan on adding other methods, as implemented and reviewed in [3] and [43].

These additions should help with both speed and accuracy of NS-DMD. In the meantime, the current rendition of NS-DMD works very well in many simulations and on a range of different empirical data, and the method already has the power to elucidate systems that were previously intractable.

## ACKNOWLEDGEMENTS

Special thanks to B. Brunton for discussions on DMD and non-stationary methods, and members of the Center for Computational Neuroscience for useful NS-DMD discussions. Special thanks to A. Husain and P. Baela for their support.

## APPENDIX A CODE AVAILABILITY

Python code implementing NS-DMD and worked examples can be found at https://github.com/learning-2-learn/nsdmd. All simulations, including supplementary ones, are included here.

## APPENDIX B HYPER-PARAMETERS

All hyperparameters are listed for the simulations, LFP data, and SST data. The parameters are listed in the order they appear in NS-DMD, and we reference the steps listed in Sec. II-B4.

- Step 1 (optional) we bandpass the data for the frequency decay simulation (Sup. Sec. S2-A) and trim 1500ms off the ends. We bandpass the LFP data (Sec. III-B) and trim 500ms off the ends.
- Step 2: all simulations use a window size of 500ms and a stride of 100ms except for the frequency drift simulation (Sec. III-A2), where we use a window size of 200ms and a stride of 50ms. All simulations use an OPT-DMD rank of 4 except for the four assembly simulation (Sec. III-A1), where we use 2, 4, and 6.
- Step 3:, the similarity thresholds are listed in Table 1.
- Step 4 (optional): when using an OPT-DMD rank of 6 in the four assembly simulation, we require two or more consecutive similar modes in a row. In the LFP and SST data, we require three or more consecutive similar windows in a row.
- Step 5: for all simulations, we smooth the frequency in **S** with a moving average of size 51ms.
- Step 5 (optional): we include a channel specific temporal lag in for the simulation in Sup. Sec. S2-B.
- Step 6 (optional): for the frequency decay simulation (Sup. Sec. S2-A), we remove all individual modes where the reconstruction error is less than 0.2.
- Step 7 (optional): In all simulations except those listed below, we feature select using the exact method (Sup. Sec. SA-1), and we use a variance threshold of 0.01. In the cases of using a large rank in Sec. III-A1 or in the frequency die off simulation (Sup. Sec. S2-A), we use gradient descent with a maximum number of iterations of 5. In all simulations, we run the SBS feature selection algorithm and terminate at 1 mode. In the frequency decay simulation, (Sup. Sec. S2-A), we use the SFS algorithm and terminate at 30 modes.
- Step 8: the parameters for gradient descent are described Table 1. In all simulations, we use a maximum iteration of 100, a learning rate of 0.01, a momentum of 0.9, and a low pass filter of the initial guess at 2Hz.

**TABLE 1:**
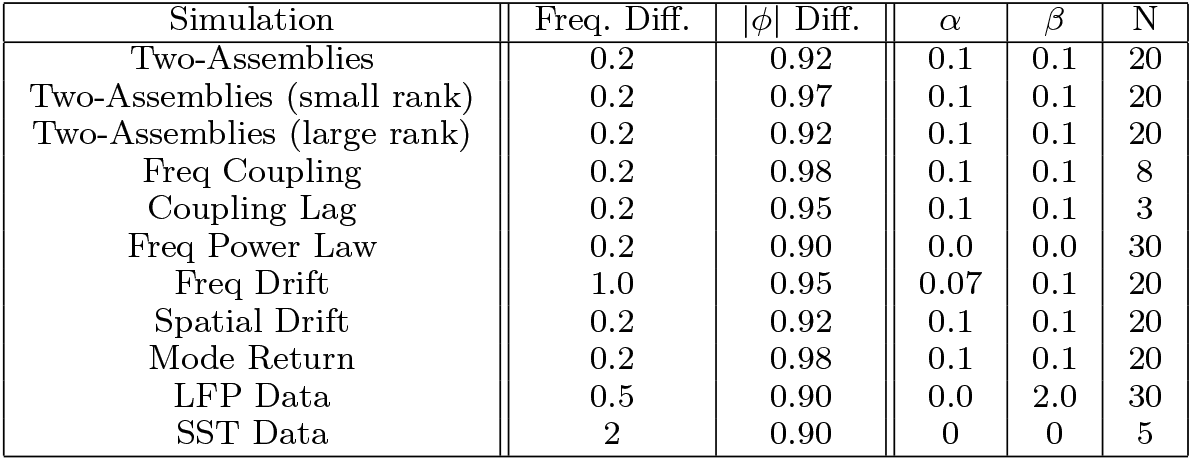
Table of similarity parameters (L), which describes the thresholds for different windows to be considered “similar”. Similar modes are grouped together. We omit the *ϕ* angle similarity threshold, which we set to 10 for all cases. This threshold is compared to the mean squared error. (R) describes the hyperparameters for gradient descent. For the *β* term and averaging between iterations, we use N consecutive time points.

| Simulation | Freq. Diff. | $\vert \phi\vert $ Diff. | $\alpha$ | $\beta$ | N |
| --- | --- | --- | --- | --- | --- |
| Two-Assemblies | 0.2 | 0.92 | 0.1 | 0.1 | 20 |
| Two-Assemblies (small rank) | 0.2 | 0.97 | 0.1 | 0.1 | 20 |
| Two-Assemblies (large rank) | 0.2 | 0.92 | 0.1 | 0.1 | 20 |
| Freq Coupling | 0.2 | 0.98 | 0.1 | 0.1 | 8 |
| Coupling Lag | 0.2 | 0.95 | 0.1 | 0.1 | 3 |
| Freq Power Law | 0.2 | 0.90 | 0.0 | 0.0 | 30 |
| Freq Drift | 1.0 | 0.95 | 0.07 | 0.1 | 20 |
| Spatial Drift | 0.2 | 0.92 | 0.1 | 0.1 | 20 |
| Mode Return | 0.2 | 0.98 | 0.1 | 0.1 | 20 |
| LFP Data | 0.5 | 0.90 | 0.0 | 2.0 | 30 |
| SST Data | 2 | 0.90 | 0 | 0 | 5 |

