## Supplemental Material for "Non-Stationary Dynamic Mode Decomposition"

Digital Object Identifier

### Non-Stationary Dynamic Mode Decomposition, Supplemental Material

In the supplemental materials, we add two extra approaches one can take with NS-DMD. These approaches aren't recommended due to additional noise or lack of additional information, but they are included for completion. We also add a variety of simulations for interesting edge cases.

#### S1. ADDITIONAL OPTIONAL STEPS

##### A. EXACT METHOD TO BYPASS GRADIENT DESCENT

By looking at NS-DMD, one could imagine directly solving for the modulation $\mathbf{F}$ instead of performing gradient descent. This approach tends to include noise in the result, but it is included here for completion. It can speed up some computations, so it can be useful during feature selection.

A direct approximation of $\mathbf{F}$ begins with the following:

$$\mathbf{x}_j \approx \sum_{\hat{k}} \hat{\mathbf{s}}_{:\hat{k}j}\hat{f}_{\hat{k}j} = \sum_{k} \mathbf{s}_{:kj} f_{kj} + \mathbf{n}_j, \tag{S1}$$

where $\mathbf{n}_j \in \mathbb{R}^N$ is a new term representing the noise. Hats $\hat{}$ indicate approximations of the true modes. Any true mode k in $\boldsymbol{\mathcal{S}}$ that we do not find in the modes $\hat{k}$ of $\hat{\boldsymbol{\mathcal{S}}}$ is included in $\mathbf{n}$. For each snapshot $t_j$ and mode k, we take the gradient of the L2 norm (Eq. 9 in the main text) and set $\hat{f}_{k'\neq\hat{k},j} \equiv 0$ to get:

$$\hat{f}_{\hat{k}j} = \frac{\mathbf{x}_j \hat{\mathbf{s}}_{:\hat{k}j}}{|\hat{\mathbf{s}}_{:\hat{k}j}|^2}. \tag{S2}$$

This effectively finds individual $\hat{\mathbf{F}}$s for each mode. Since our goal is to estimate $\mathbf{F}$, not $\hat{\mathbf{F}}$, we substitute the true solutions for $\mathbf{X}$

$$\hat{f}_{\hat{k}j} = \frac{(\mathbf{S}_j \mathbf{f}_j + \mathbf{n}_j)\hat{\mathbf{s}}_{:\hat{k}j}}{|\hat{\mathbf{s}}_{:\hat{k}j}|^2}, \tag{S3}$$

and define the following terms:

$$b_{k\hat{k}j} \equiv \frac{\mathbf{s}_{:kj}\hat{\mathbf{s}}_{:kj}}{|\hat{\mathbf{s}}_{:\hat{k}j}|^2} \quad \text{and} \quad \tilde{n}_{\hat{k}j} \equiv \frac{\mathbf{n}_j \hat{\mathbf{s}}_{:\hat{k}j}}{|\hat{\mathbf{s}}_{:\hat{k}j}|^2}. \tag{S4}$$

Our equation reduces to

$$\hat{\mathbf{f}}_j = \mathbf{f}_j \mathbf{B}_j + \tilde{\mathbf{n}}_j. \tag{S5}$$

The next few steps serve to invert the above equation to find $\mathbf{F}$.

For the inversion, we approximate $\hat{\boldsymbol{\mathcal{S}}} \approx \boldsymbol{\mathcal{S}}$ and remove $\mathbf{n}_j$. In other words, we assume that we have captured all relevant modes in $\hat{\boldsymbol{\mathcal{S}}}$, and we ignore any additional modes.

$$\hat{\mathbf{f}}_j \approx \mathbf{B}_j \mathbf{f}_j, \tag{S6}$$

where the rank of $\mathbf{B}_j$ is less than its number of rows/columns. We use an SVD [1] approach here:

$$\mathbf{U}_j \tilde{\mathbf{B}}_j \mathbf{U}_j^{\dagger} \mathbf{f}_j = \hat{\mathbf{f}}_j, \tag{S7}$$

where $\tilde{\mathbf{B}}_j$ is diagonal and $\mathbf{U}_j$ is a unitary matrix that diagonalizes $\mathbf{B}_j$. We reduce $\tilde{\mathbf{B}}_j$ to a low rank r by only including large variance terms, since $\tilde{\mathbf{B}}_j$ is linearly dependent. We use a variance threshold of 0.01. We invert and find an estimate of $\mathbf{F}$:

$$\mathbf{f}_j \approx \mathbf{U}_j \tilde{\mathbf{B}}_j^{-1} \mathbf{U}_j^{\dagger} \hat{\mathbf{f}}_j. \tag{S8}$$

The steps to find $\mathbf{F}$ are then to calculate $\hat{\mathbf{f}}$ in Eq. S2, $\boldsymbol{\mathcal{B}}$ in Eq. S4, and find the SVD of $\boldsymbol{\mathcal{B}}$.

##### B. INDIVIDUAL GLOBAL MODULATION PER CHANNEL

It may also be possible that the time dependent functions $\mathbf{f}(t)$ are different for each recording channel. Specifically, we analyze systems such that $x_i(t) \sim x_i(t + \Delta t_{ij})$, where the time dependence of channels i and j are offset by $\Delta t_{ij}$. In these systems, both the phase of the spatiotemporal modes $\boldsymbol{\mathcal{S}}$ and the modulations $\mathbf{F}$ are related by the same temporal offset $\Delta t_i$. We can define the time dependent function $\mathbf{g}_i(t) \equiv f(t + \Delta t_i)$ to distinguish the time dependence of every channel from the global modulation $f(t)$. The spatiotemporal modes are the same as before: $\mathbf{S}(t) \equiv \boldsymbol{\Phi}(t)e^{i\angle\boldsymbol{\Lambda}(t)t}$, where the phase of each channel $\angle\boldsymbol{\Phi}_i = \angle\boldsymbol{\Lambda}(t)\Delta t_i$. We then have:

$$\mathbf{x}_i = \sum_{k} \mathbf{s}_{:ki}\mathbf{g}_{:ki}. \tag{S9}$$

To calculate the temporal lag, $\Delta t_i$, one can use $\angle\phi$. However, this only works when the lag is less than half of a period; when any larger, the total time delay is ambiguous due to periodicity. A more general method is to instead check the reconstruction error for all possible temporal lags. Each lag can be compared to knowledge of the system: e.g delays between measurements in fluid flow or expected transit times between neurons in the brain. We expect the lag to be approximately a $2\pi$ multiple of the phase of $\phi$. However, without knowledge of the system, we have not found a general way to automate finding the exact temporal lag.

From simulations in Sec. S2-B, we find that this additional feature does not help improve the fit enough to warrant its use. It is included as an optional step.

#### S2. ADDITIONAL SIMULATIONS

We now apply NS-DMD to more simulated data, in situations that either require optional steps in the NS-DMD method or are specifically interesting to some communities. We simulate synthetic data with the following generic equation:

$$\mathbf{x}(t_i) = \sum_k f_k(t_i)\mathbf{g}_k(t_i + \frac{\phi_{P,k}(t_i)}{\omega_k(t_i)})\phi_{A,k}(t_i) * \cos(\sum_{j=0}^{i-1} \omega_k(t_j)\Delta t + \phi_{P,k}(t_i)), \tag{S10}$$

##### A. POWER LAW SPECTRAL DISTRIBUTION

An important issue is having a system composed of many different frequencies at different magnitudes of power. One example, that occurs in many communities, such as physics biology, economics, etc. [5], are frequency power laws. It is common for signal to occur where the power fluctuates like $1/f^{\alpha}$. This case requires the optional step of bandpassing the data into smaller bands to find modes with smaller amplitudes.

We simulate power law decay with random spatial modes $\mathbf{\Phi}$ that occur in 5Hz increments from 2Hz to 100Hz, where the amplitude occurs with an exponent of $\alpha = -2$. For simplicity, we do not include any non-stationarity in the amplitude, which corresponds to the following simplification of Eq. S10:

$$\mathbf{x}(t_i) = \phi_{A,k}\cos(\omega_k t_i + \phi_{P,k}). \tag{S11}$$

Random noise is added to each frequency with a standard deviation of 0.5. Fig. S1A shows the power spectrum of an example channel. Due to pink noise, we find an exponent of $-3.4$ when fitting a line in log space. Modes become blended together at higher frequencies. I.e. for a 1s increment, 2 and 3Hz are very different while 92 and 93Hz are very similar. Thus, this simulation tests NS-DMD at the ability to segregate small and large frequency differences.

The result of NS-DMD is a relatively flat spectrum of $\mathbf{F}$ (Fig. S1B), exactly under the simulation parameters. Note that we normalize $\mathbf{F}$ for every mode for visualization. We show the mean of $\mathbf{F}$ for all frequencies in Fig. S1C, where we see that the amplitude decays like the true power law.

NS-DMD doesn't perform perfectly at higher frequencies, where in this example there is an extra 67Hz mode and lack of 87Hz mode. NS-DMD tends to perform better at lower frequencies where the modes are better segregated from each other.

##### B. FREQUENCY COUPLING AND INDEPENDENT CHANNEL MODULATION

We turn our attention to a neuroscience issue, where it's believed that low frequencies modulate high frequencies via frequency coupling [2, 4, 3, 6]. The lower frequency signal likely has a signal with an unknown exact form. Thus, we simulate a square wave, where all possible frequencies are part of the low frequency signal. The square wave switches between 0 and 1 with a periodic frequency of 5Hz (Fig. S2). Using NS-DMD, we find the dominating 5Hz frequency in Fig. S2. Since NS-DMD works with a square wave, it likely works with any periodic function.

We modify this simulation to explore the case when individual channels are independently modulated, as developed in Sup. Sec. S1-B. For this example, each channel has the same modulation, offset by a temporal phase, which corresponds to the following simplification of Eq. S10:

$$\mathbf{x}(t_i) = \mathbf{g}_k(t_i + \frac{\phi_{P,k}}{\omega_k})\phi_{A,k}\cos(\omega_k t_i + \phi_{P,k}). \tag{S12}$$

We generate the data with two assemblies. The first assembly has a constant amplitude at 30Hz. For the second assembly, each channel is modulated by a square wave that stops around $t = 1000$ms (Fig. S3B).

While running NS-DMD, we implement the optional step of including temporal lags for each channel based on the angle of $\mathbf{\Phi}$. In Figs. S3A and S3B, we show that NS-DMD recovers the correct global modulation $\boldsymbol{\mathcal{G}}$ for each individual channel. We also ran NS-DMD without the optional temporal lags, where NS-DMD finds an approximation of $\boldsymbol{\mathcal{G}}$ (Fig. S3C). With this in mind, along with the requirement that the temporal delay is less than half a period, the optional inclusion of temporal lags in NS-DMD may not be needed.

##### C. MODES RETURNING WITH NEW PHASE

When $\mathbf{F} = 0$, the phase is undefined, and it's possible that an assembly will turn off and back on with a different phase. If the phase difference is small, we expect NS-DMD to find one mode that explains the assembly before and after $\mathbf{F} = 0$. If the phase difference is large, up to $\pi$, we expect NS-DMD to find two modes that explain the data. To analyze the behavior of NS-DMD on this effect, we generate synthetic data from two assemblies. The first mode has a constant $f(t)$ for the duration of the dataset at 12Hz with a randomly generated spatial mode $\phi$. The second assembly turns off at $\sim$ 500ms and back on at $\sim$ 1000ms with a frequency of 5Hz. The shape of $f(t)$ is shown in Fig. S4. The spatial mode $\phi$ is randomly generated, and the phase $\angle\phi$ of each channel is advanced 0, $\pi/2$, and $\pi$ radians for Figs. S4A,D, Figs. S4B,E, and Figs. S4C,F respectively when the mode returns at $\sim$ 1000ms.

To analyze the behavior of NS-DMD, we run NS-DMD on each case with a final subset of either two Figs. S4A-C or three modes Figs. S4D-F. As expected, when the phase does not change, we find two modes is sufficient to capture both assemblies Fig. S4A. If instead, we use

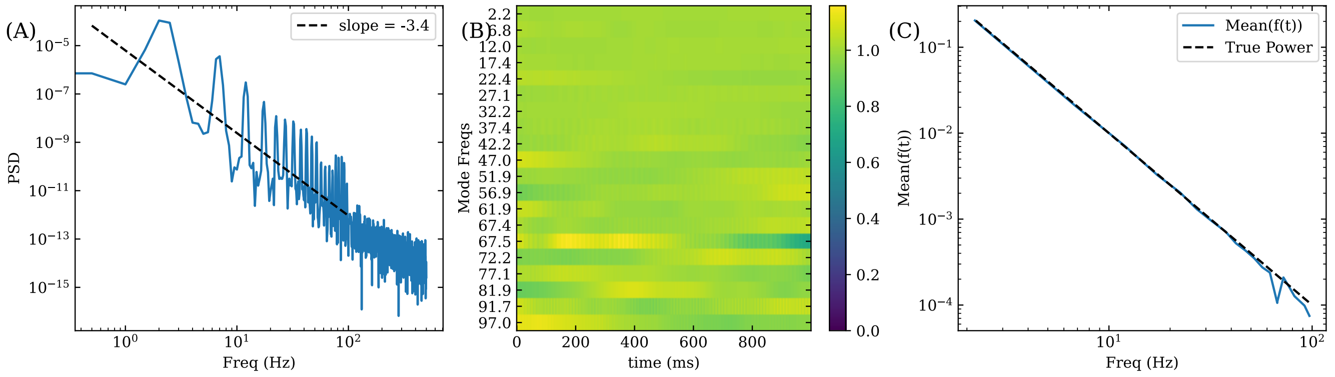


FIGURE S1: (A) shows the Power Spectral Density (PSD) of the first channel. The peaks indicate the frequencies of each mode, which have a power law decay of $-2$. The frequencies occur at $2 + 5n$Hz. The background noise is added at a slope of $-2.5$. All channels have a similar PSD. (B) shows $f(t)$ for each mode, normalized by their mean. NS-DMD approximately recovers the true, flat distribution equal to 1. However, there is an extra 67Hz mode and no 87Hz mode. (C) shows the true power law decay (dotted line) compared to the mean of $f(t)$ of each mode. NS-DMD recovers the true power law.

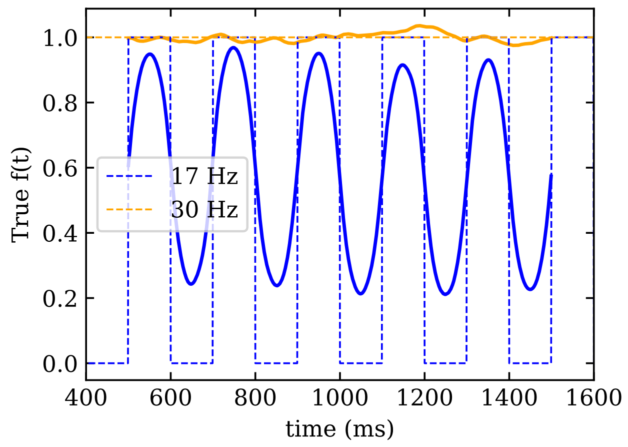


FIGURE S2: Comparison of the true (dotted lines) and fitted $f(t)$ (solid lines). NS-DMD is able to find the dominate, 5Hz Fourier mode of the this coupling. Depending on the amount of smoothing, one can be more or less accurate during the steepest portions of the square wave. Success with NS-DMD on square wave coupling demonstrates the ability to work on any periodic coupling. This is due to the Fourier series of a square wave including all frequencies.

three modes, we find a mixture of two modes to explain the leaving and returning assembly Fig. S4D.

In the other simple case where the phase is advanced $\pi$ radians, we find that two modes only recovers the assembly at the beginning of the time-series (Fig. S4C). Three modes fully recovers the data (Fig. S4F).

We see a mixture of effects in the case where we advance the phase $\pi/2$ radians. When converging to two modes, NS-DMD partially recovers the assembly when it returns. With three modes, NS-DMD recovers $\mathbf{F}$, and two of the modes mix a little bit.

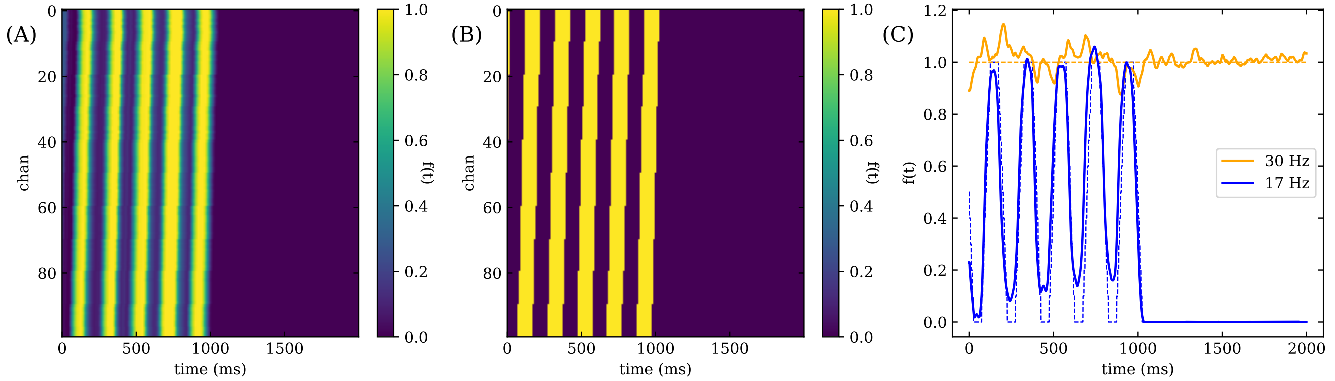


FIGURE S3: We show the results of independent modulations for each channel at 17Hz. We compare the fit f(t) for each channel in (A) to the true f(t) in (B), where we find that NS-DMD recovers the ground truth. Note that this only works when the overall lag is less than a period at the modes frequency. We also show in (C) a comparison of the true f(t) (dotted lines) and fitted f(t) (solid lines) when we do not include a temporal lag for **F**. NS-DMD still recovers the correct overall global modulation. The true 17Hz modulation is trapezoidal, which is due to averaging a square wave with different temporal lags.

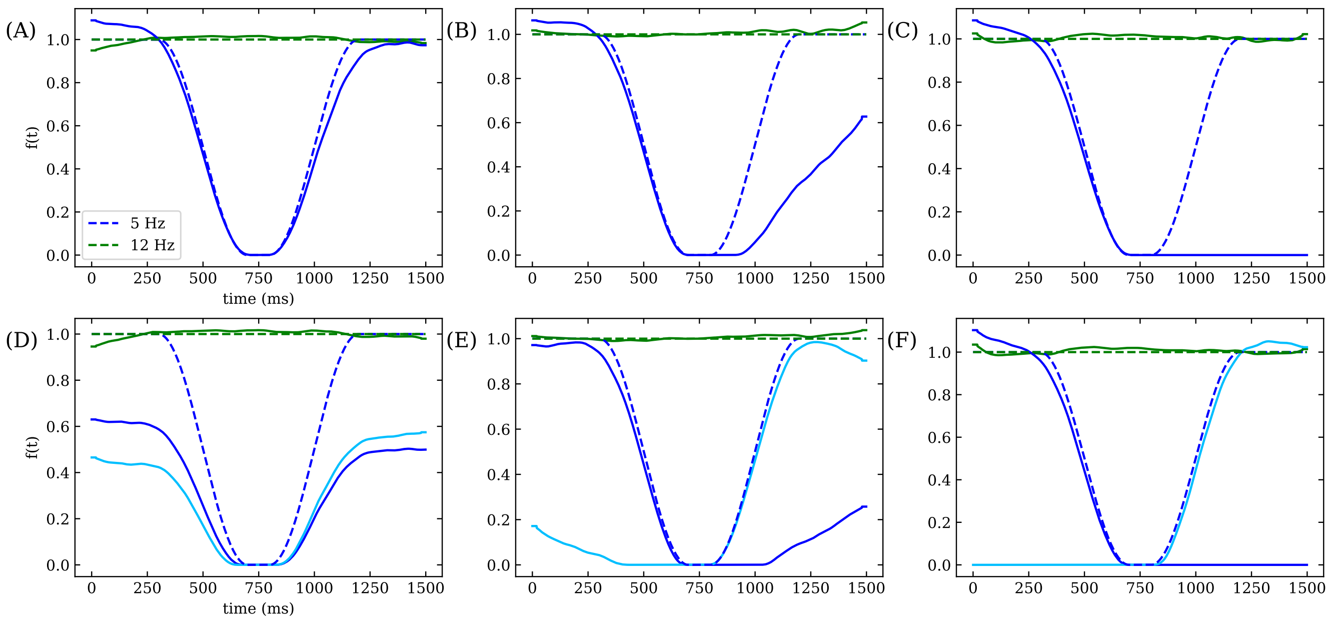


FIGURE S4: This figure shows the true f(t) (dotted lines) and approximated f(t) (solid lines) for different cases of the returning assembly simulation. There are two assemblies, one with a constant amplitude at 12Hz, and one that turns off and back on at 5Hz. The spatial mode $\phi$ is random for each assembly, and the overall phase of $\phi$ is advanced 0, $\pi/2$, and $\pi$ radians for (A)/(D), (B)/(E), and (C)/(F) respectively. In (A)-(C), we converge to two modes with NS-DMD. In (D)-(F), we converge to three modes with NS-DMD. When the phase doesn't change, we find that two modes best fit the data (A). When the phase advances $\pi$ radians, we find that three modes best fit the data. Anywhere in between, we find that two modes underfits the data, and three modes can show a mixing effect.
